# Structural characterisation of the complete cycle of sliding clamp loading in *Escherichia coli*

**DOI:** 10.1101/2023.07.20.549978

**Authors:** Zhi-Qiang Xu, Slobodan Jergic, Allen T.Y. Lo, Alok C. Pradhan, Simon H.J. Brown, James C. Bouwer, Harshad Ghodke, Peter J. Lewis, Gökhan Tolun, Aaron J. Oakley, Nicholas E. Dixon

## Abstract

Ring-shaped DNA sliding clamps are essential for DNA replication and genome maintenance. Clamps need to be opened or trapped open and chaperoned onto DNA by clamp loader complexes (CLCs). Detailed understanding of the mechanisms by which CLCs open and place clamps around DNA remains limited. Here, we present a series of six structures of the *Escherichia coli* CLC bound to an open or closed clamp on and off a primer-template DNA that represent all intermediates in the clamp loading process. We show that the ATP-bound CLC first binds to a clamp, then constricts to hold onto it. The CLC then expands to open the clamp with a gap large enough for double-stranded DNA to enter. Upon binding to DNA, the CLC constricts slightly, allowing ATP hydrolysis and clamp closing around DNA. Although both yeast and *E. coli* CLCs open clamps by crab claw-like motions, they do it by the CLC expanding in opposite directions. These structures provide critical high-resolution snapshots of clamp loading by the *E. coli* CLC, revealing how the molecular machine works.

## Introduction

Rapid and processive replication of chromosomal DNA requires DNA polymerases to be tethered to primer-template (p/t) DNA by a sliding clamp that encircles and slides on double-stranded (ds)DNA^1–4^. Sliding clamps also serve as a tether for many other proteins involved in DNA metabolism^5^. These ring-shaped clamps^6–8^ need to be opened and chaperoned onto a p/t DNA junction in an ATP-dependent process by a multisubunit clamp loader complex (CLC)^2,9^. Frequent and rapid loading of sliding clamps onto newly-primed sites by the CLC is critical for timely production of Okazaki fragments during lagging-strand DNA synthesis^10^.

Structures and functions of both clamps and CLCs from different organisms are conserved^3,4^. All sliding clamps contain six structurally-similar domains (Fig. 1a) and CLCs are complexes comprised of five AAA+ proteins (ATPases associated with various cellular activities)^11^ (Fig. 1b). The *Escherichia coli* sliding clamp contains two β subunits arranged as a head-to-tail dimer^6^, while the clamp from bacteriophage T4 and eukaryotic and archaeal PCNAs (proliferating cell nuclear antigens) are trimers^3,4^. While the *E. coli* and eukaryotic clamps are stable closed rings, the T4 clamp can open spontaneously^12,13^. The *E. coli* CLC is comprised of five core subunits δτ_n_γ_3–n_δ’ in positions A–E, respectively (Fig. 1b), and two accessory proteins, ψ and χ^10^, whereas eukaryotic CLCs (RFC, replication factor C) contain five different proteins Rfc1−5^3,4^, and the T4 CLC contains gp62 (in position A) and four gp44 proteins. The *E. coli* τ subunit is a longer version of γ with two additional C-terminal domains that interact with the DnaB helicase and the Pol III DNA polymerase α subunit, serving as an organisational hub in the Pol III holoenzyme^10^ (Fig. 1c). Each CLC protein consists of three domains. The N-terminal RecA-type ATPase domain (Domain I) and the middle lid domain (II) form a AAA+ module. The C-terminal domains (III) form a disc-like collar that caps and stabilises the AAA+ domain^3,4^. Interactions of the CLC proteins with clamps are primarily mediated by a peptide with a consensus sequence called the clamp-binding motif (CBM) binding to surface crevices on the clamps^14,15^.

**Fig. 1.**
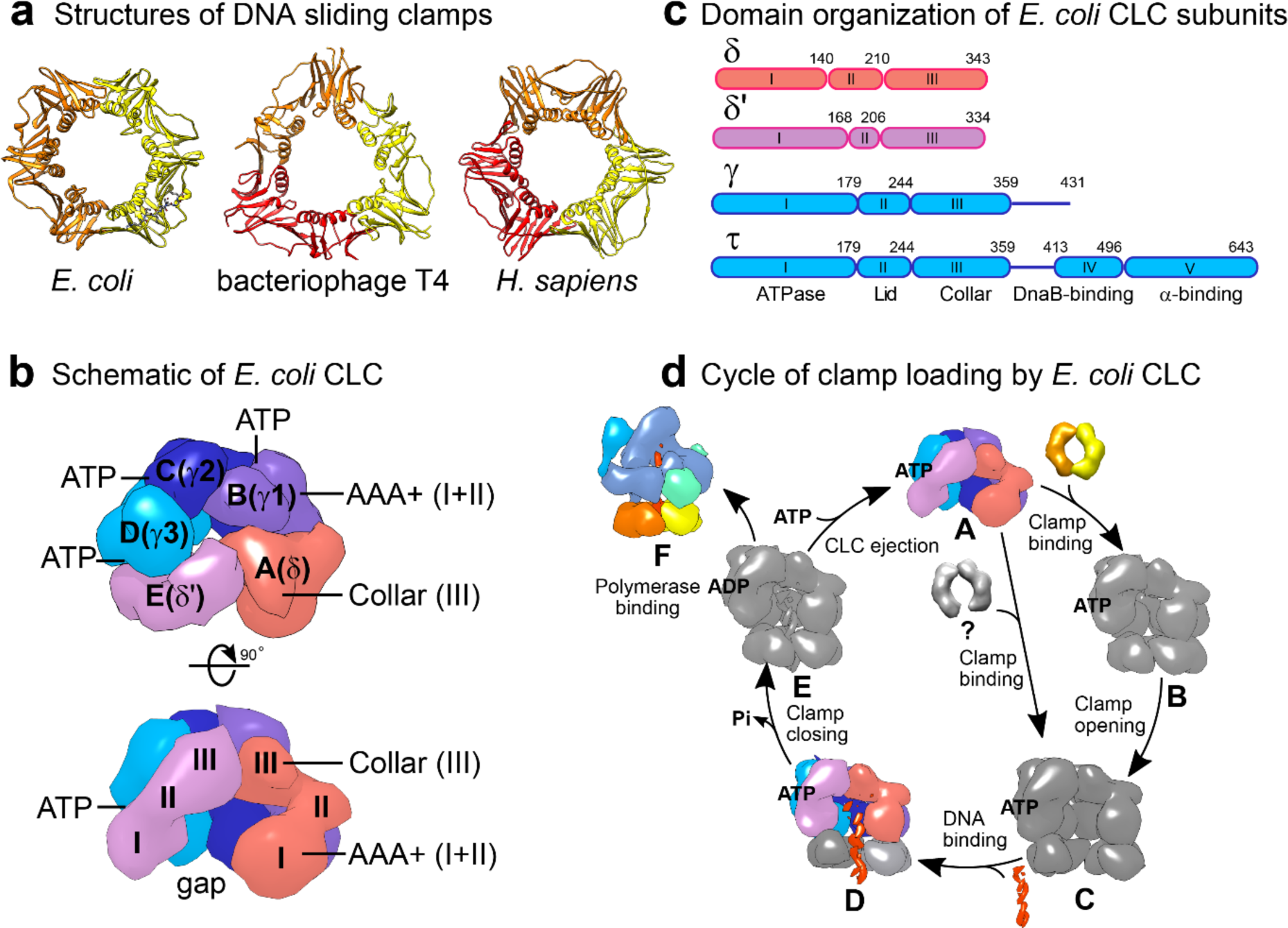
The clamp loading process and its participants. **a**, Sliding clamps from *E. coli* (PDB 1MMI^41^), phage T4 (1CZD^8^) and humans (1VYJ^57^) are closed rings containing six structurally similar domains. The *E. coli* clamp is dimeric, while T4 and eukaryotic clamps are trimers. **b**, Views of the CLC from collar to AAA+ domain (top view, top panel) and front to back (side view, bottom panel). The *E. coli* CLC core subunits, δ, γ1, γ2, γ3 and δ’ (coloured salmon, medium pink, medium blue, deep sky blue and plum, respectively), are arranged anticlockwise in canonical positions A−E. ATP molecules (or analogues) are bound between Domains I and II of γ subunits at the γ1γ2, γ2γ3 and γ3δ’ interfaces. **c**, Domain organisation of *E. coli* CLC core subunits. All subunits contain AAA+ ATPase domains (I and II) and a Collar domain (III). The τ subunits contain an additional Domain IV that interacts with the DnaB helicase and Domain V that binds tightly to the Pol III α subunit. **d**, Schematic of clamp loading in *E. coli*. To load a clamp, the ATP-bound CLC (**A**) first binds to a β_2_ clamp (**B**)^4,20–22^ and opens it (**C**)^22–25^. Alternatively, the β_2_ clamp may open spontaneously that is unlikely and the CLC trap it in an open state (**C**). Then primer-template (p/t) DNA passes through the gap of the clamp and binds in the chamber of the CLC (**D**)^27^ either *via* its ssDNA or dsDNA portion depending on the size of the gap in β_2_. Upon binding and recognition of a p/t DNA junction, ATP hydrolysis occurs as the β_2_ clamp closes on DNA (**E**)^28,29^ and the CLC is ejected leaving the loaded clamp on p/t DNA for binding of the Pol III αεθ core to the clamp (**F**)^3,4^. Structures already reported for the *E. coli* CLC are in colours (**A** and **D** without β_2_); structures only available for other organisms are in grey (**B**, yeast and human; **C**, yeast; **D** and **E**, phage T4 and yeast).

Three ATP molecules bind at AAA+ subunit interfaces involving the three γ/τ subunits. In the absence of ATP, the AAA+ domains of the *E. coli* CLC are not well-organised^16,17^ and incapable of high-affinity clamp binding^4,18^. ATP binding induces conformational changes in the CLC that enable clamp binding^19^. To load a clamp (Fig. 1d), the ATP-bound CLC (state **A**) either binds to a clamp (**B**)^4,20–22^ and opens it (**C**)^19,22–25^ or traps a spontaneously opened clamp (**C**)^26^. The CLC-open clamp complex can directly bind to p/t DNA if the gap in the clamp is large enough for dsDNA to pass. Otherwise, the complex has to use a “filter and slide” mechanism in which only ssDNA can pass through the gap and p/t DNA has to slide up the chamber of the CLC (**D**)^27,28^. Upon binding and recognition of a p/t DNA junction, the clamp closes on DNA (**E**)^28,29^, followed by ejection of the CLC and binding of a DNA Pol III core to the clamp (**F**)^3,4^. The timing of hydrolysis of the three ATP molecules is uncertain. It seems likely that clamp closing and CLC ejection is a concerted process during which all three ATPs are hydrolysed rapidly and sequentially. Careful correlation of the pre-steady state kinetics of ATP hydrolysis with p/t DNA and clamp binding and release suggest the hydrolysis of two ATPs occurs during clamp closing and one during clamp ejection^29–32^. However, which of the ATPs are hydrolysed during each of these steps is unknown.

High-resolution structures of CLC-clamp complexes are important to understand how CLCs load sliding clamps onto DNA. For the much-studied *E. coli* CLC, only structures of free CLC^16,17^ (**A**) and CLC bound to a p/t DNA (**D**) but without a β_2_ clamp^27^ are available (coloured structures in Fig. 1d). While related structures of CLC-clamp complexes from phage T4^28^, yeast^20,33–36^ and human^21^ have provided snapshots of clamp loading intermediates for those systems, questions remain for the *E. coli* CLC: 1. Does it actively open the clamp^22,25^ or trap an open clamp^26^? 2. If it actively opens a clamp, how does it achieve that? 3. Is the gap on an open β_2_ clamp large enough to allow direct passage of dsDNA or only ssDNA? 4. When is ATP hydrolysed and at which sites?

Here, we report a series of high-resolution cryogenic electron microscopy (cryo-EM) structures of *E. coli* CLC•β_2_ complexes on and off p/t DNA that provide a comprehensive view of the clamp loading cycle. Our studies reveal that the CLC first binds to β_2_ *via* the CBM of δ, then constricts to hold on to β_2_ through its simultaneous binding to subunits δ, γ1, γ3, and δ’. The CLC then actively opens β_2_ by crab claw-like motions to produce a gap large enough for direct passage of dsDNA. Surprisingly, the *E. co*li CLC opens β_2_ by moving the AAA+ modules of subunits A (δ) and B (γ1) outwards anticlockwise in hinge-like motions that are different from the yeast RFC that moves AAA+ modules C, D and E outwards clockwise. Thus, despite conserved structures and functions, the mechanisms of clamp opening by CLCs of *E. coli* and *Saccharomyces cerevisiae* have diverged.

## Results

### The *E. coli* CLC binds, constricts and holds onto a closed β_2_ clamp before opening it

The β_2_ clamp is a very stable homodimer^12,13,37,38^ that needs to be opened for loading onto a p/t DNA by the CLC^4^. To understand how the CLC binds and opens the β_2_ clamp, we first used surface plasmon resonance (SPR) to test how strongly the CLC and β_2_ interact in the presence of ATP, ATPγS – an ATP analogue poorly hydrolysed by the CLC^19,39^, or ADP•AlF_x40_ – a non-hydrolysable ATP transition-state mimic. The full seven-subunit δτ_3_δ’ψχ CLC was first bound to streptavidin on the SPR chip surface *via* the (biotinylated) χ subunit, then serially diluted β_2_ solutions were flowed over CLC to interact. We found that β_2_ bound to the CLC strongly with ATP and ATPγS (*K*_D_ of 2.7 ± 0.3 and 13.3 ± 0.1 nM, respectively), while binding was weaker with ADP•AlF_x_ (400 ± 15 nM) (Extended Data Fig. 1). Therefore, we chose to use ATPγS as nucleotide and solved the structures of δγ_3_δ’ψχ bound to β_2_ by single-particle cryo-EM using a sample pre-treated with the amine-reactive crosslinker bis(sulfosuccinimidyl)suberate (BS^3^)^21,33^ to further stabilise the complex.

The data reveal a mixture of CLC•β_2_ structures with open and closed β_2_ clamps (Extended Data Table 1 and Fig. 2, 3), suggesting that ATPγS binding may not simply open the clamp, but produce an equilibrium mixture of open and closed states. Reconstructions of the CLC bound either to the closed (CLC•β_2_^closed^, refined to 3.7 Å resolution) (Fig. 2a) or to the open β_2_ clamp (CLC•β_2_^open^, 2.7 Å) (Fig. 3a) were made, and the structures of CLC•β_2_^closed^ and CLC•β_2_^open^ were then built using crystal structures of the *E. coli* CLC^27^ and β_241_. In both maps, the whole of the δ and δ’ subunits and Domains I−III of the γ subunits are resolved; flexible regions beyond Domain III of γ subunits (Fig. 1c) were not visible in the density maps. There is weak density of ψ beyond its well-resolved N-terminal γ-binding peptide (Fig. 3a) that binds across the collars of all three γ subunits as revealed previously (PDB 3GLI)^27^, allowing placement of the ψχ complex using its reported crystal structure (PDB 1EM8)^42^. Density interpreted as ATPγS is present at all three ATP-binding sites on the CLC at the γ1γ2, γ2γ3 and γ3δ’ interfaces in both structures (Extended Data Fig. 2d). There is also unassigned density ascribed to a short peptide nestling in a shallow groove on Domain I of δ’, next to its N-terminus and potentially stacking with the aromatic ring of Trp3 (Fig. 2a, 3a). This is likely a part of the flexible C-terminal region of one of the γ subunits; the role of this interaction is unknown.

**Fig. 2.**
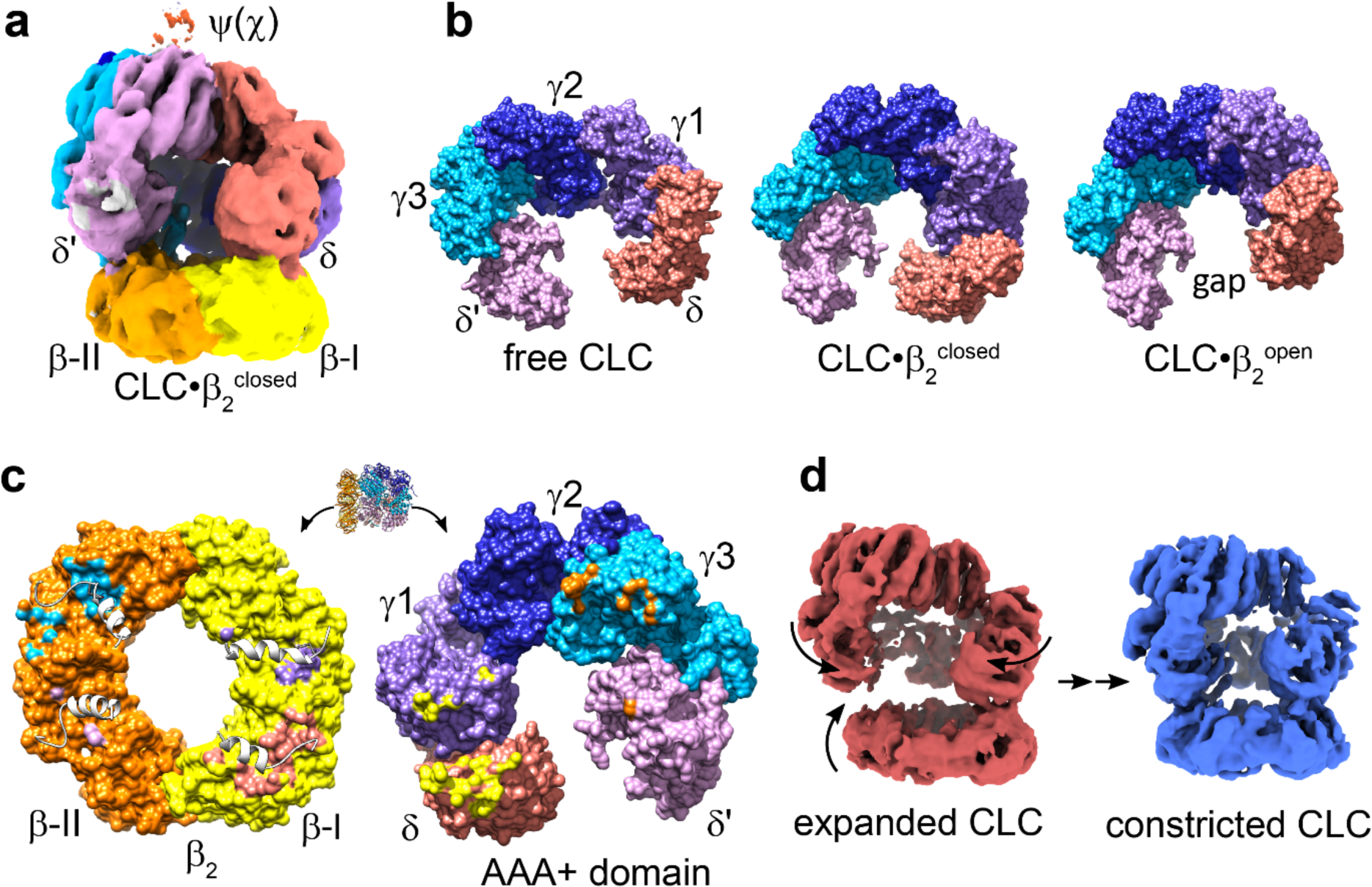
Clamp binding by the *E. coli* CLC is highly dynamic. **a**, Cryo-EM map of *E. coli* CLC bound to a closed β_2_ clamp (CLC•β_2_^closed^) in the absence of p/t DNA. Cryo-EM densities corresponding to the δ, γ1, γ2, γ3, δ’ and ψ subunits of the CLC are coloured salmon, medium purple, medium blue, deep sky blue, plum and orange red, respectively. The β-I subunit of β_2_ is coloured yellow and β-II orange. Density next to Domain I of δ’, presumed to be from the unstructured C-terminal region of a γ/τ subunit is coloured grey (see also Fig. 3a). The colouring scheme is kept throughout unless stated otherwise. **b**, The AAA+ modules (shown without the Collar domain) of γ1 and δ in CLC•β_2_^closed^ encroach into the gap between δ and δ’. Structures are aligned on γ2, which is barely perturbed among aligned structures. The AAA+ domains of the free δγ_3_δ’ complex (PDB 1XXH)^17^ are disorganised, while those of CLC•β_2_^closed^ and CLC•β_2_^open^ are more regularly packed. **c**, CLCβ interfaces on β_2_ (left) and CLC (right) of CLC•β_2_^closed^. CLCβ interactions are mediated by peptides of the CLC subunits that are located on an α-helix and the following loop in their Domains I, which are presented as white ribbons on the surfaces of β_2_. β_2_ and the CLC are presented side-by-side by separating and turning them 90° in opposite directions. Buried areas are coloured as for their interacting counterparts; those coloured in salmon (δ) and deep sky blue (γ3) are in the two symmetry-related canonical peptide binding sites of β_2_. **d**, Dynamics of clamp binding revealed by 3D variability analysis (3DVA). Volumes of the first (brick red) and last (blue) frames of principal component_001 of 3DVA that show the most significant and biologically relevant conformational changes are presented.

**Fig. 3.**
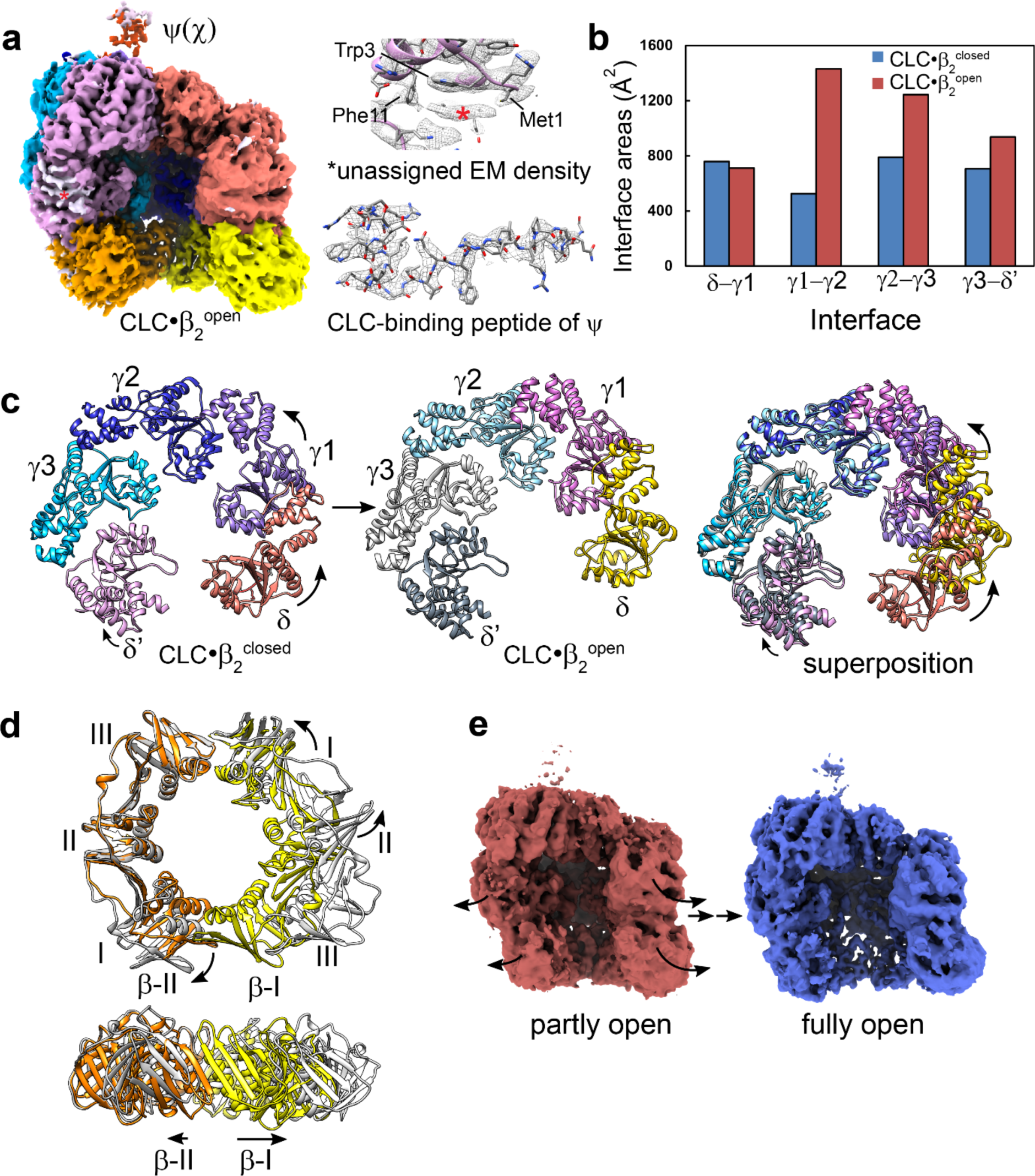
The *E. coli* CLC opens β_2_ by crab claw-like motions. **a**, Cryo-EM map of CLC bound to an open clamp (CLC•β_2_^open^) coloured as in Fig. 2. Top right, unassigned EM density near the N-terminus of δ’ presumed to be from a peptide near the C-terminus of a γ subunit (Fig. 1c) is indicated by a red asterisk. A moving window of 8-residue peptides selected from the C-terminal flexible region of γ were docked into δ’ and scored against the density. The best scoring peptide corresponded to the moderately-conserved γ residues Val403−Thr409. This assignment is yet to be verified experimentally. Bottom right, unsharpened EM density of the CLC-binding peptide of ψ (Thr2–Glu28) that bridges all three γ subunits^27^. **b**, Interface areas between AAA+ modules increase significantly in the transition from CLC•β_2_^closed^ to CLC•β_2_^open^. **c**, Conformational changes of the AAA+ modules during transition from CLC•β_2_^closed^ to CLC•β_2_^open^. While the AAA+ modules of γ2 and γ3 barely move, those of γ1 and δ and to a lesser extent δ’ swing outwards, resulting in expansion of the CLC and opening of β_2_. Structures were superimposed by aligning the γ2 subunit using MatchMaker in Chimera^58^. Arrows indicate movements of individual subunits. CLC•β_2_^open^ is coloured differently from the normal scheme for comparison. **d**, Orthogonal views of conformational changes in β_2_ during transition from CLC•β_2_^closed^ (yellow and orange) to CLC•β_2_^open^ (grey). The clamp remains mostly flat after opening. Most obvious motions are on the β-I subunit with Domain I rotating around Domain III of β-II and Domains II–III around Domain I. **e**, Conformational changes in the final stage of clamp opening revealed by 3D variability analysis. Volumes of the first and last frames of principal component_000 of 3DVA that illustrate the most significant and biologically relevant conformational changes are presented.

The *E. coli* CLC•β_2_^closed^ structure represents the first stage of clamp loading (Fig. 1d, state **B**). Compared to the free “inactive” δγ_3_δ’ complexes, whose AAA+ domains are loosely packed and not arranged to allow simultaneous binding of multiple CLC subunits to β_216,17_, the AAA+ domains (Fig. 2b) of the CLC•β_2_^closed^ complex are more regularly packed and roughly match the pseudo-6-fold symmetry of β_2_ (Fig. 2c). In this closed conformation, the CLC holds onto β_2_ by simultaneous binding of the δ, γ1, γ3 and δ’ subunits. The interactions are mediated mainly by δ and γ3 and to a lesser extent by γ1 and δ’. The δ subunit makes extensive contacts with β_2_ as seen in the crystal structure of δβ (PDB 1JQJ)^15^. The interactions are mediated by the CBM motif 65-FSLC<u>QAMSLF</u> (canonical CBM underlined) binding in the canonical peptide-binding site of β_2_, burying an interface area of 717 Å^2^. CBM residues δLeu73 and δPhe74 play an important role in the interactions by forming a hydrophobic plug that wedges into a hydrophobic pocket on the surface of β_2_. The γ3 subunit occupies the canonical peptide-binding site in the other β subunit mostly *via* binding of peptide 109-DNVQYAPAR, burying 479 Å^2^. Although three distinct reconstructions of the yeast RFC•PCNA^closed^ complex were made^33^, they all contain a PCNA hanging on one side of RFC and none shows RFC held stably onto PCNA. Therefore, the CLC•β_2_^closed^ structure represents the first step in clamp binding in which the CLC has maximised contact with β_2_ and is about to open it.

However, the AAA+ domains of CLC•β_2_^closed^ are not yet symmetrically packed. The AAA+ modules of δ and γ1 are displaced from “ideal” symmetrically-arranged positions, encroaching into the gap between δ and δ’, a space occupied by the sixth subunit in hexameric ring-shaped AAA+ proteins (Fig. 2b). As a result, only one of the arginine fingers from the γ2 (R169), γ3 (R169) and δ’ (R158) subunits that are vital to ATP hydrolysis is close enough to interact with the γ-(thio)phosphate of ATPγS (*i.e.*, δ’R158 at the γ3–δ’ interface) and the β- and γ-phosphates are poorly resolved at this resolution (Extended Data Table 2 and Extended Data Fig. 2d, left). Therefore, the closed CLC complex is in an “auto-inhibited” state, similar to the *E. coli* δγ_3_δ’^16,17^ and eukaryotic RFC•PCNA^closed^ complexes^21,33^.

On the other hand, the ATPγS binding sites in the CLC•β_2_^open^ structure (especially at the γ1γ2 and γ2γ3 interfaces) are much better organised as would be expected since ATP binding promotes clamp opening; all three arginine fingers are close enough to the γ-thiophosphate of ATPγS to participate in ATP binding (Extended Data Fig. 2d, right). Note however that the CLC•β_2_ complex has only weak ATPase activity in the absence of p/t DNA^39^, so none of the ATP-binding sites is yet poised for efficient ATP hydrolysis.

We considered that the consensus reconstruction of CLC•β_2_^closed^ may contain a mixture of similar conformations, which may explain why the refinement stalled at ∼3.7 Å resolution despite there being more particles than for our other higher-resolution reconstructions (Extended Data Fig. 2). To probe these conformations, we used 3D variability analysis (3DVA)^43^ of the refined particles. The results show that the consensus reconstruction indeed contains an ensemble of continuously changing and significantly different conformations (Fig. 2d). The process of clamp binding is thus highly dynamic with a wide range of different motions of the CLC. The most significant and biologically relevant insights gained from this analysis are: (i) δ must be the first subunit to engage in β_2_ binding. (ii) The CLC then gradually constricts to allow γ1, γ3 and δ’ to interact simultaneously with β_2_. (iii) Meanwhile, β_2_ and the CLC also move gradually closer as their interactions become stronger until the CLC holds stably onto β_2_. While the conformation of the yeast RFC•PCNA^closed^ complex also changed continuously with PCNA swinging against RFC, RFC seems to be more stable than the *E. coli* CLC and remains constricted during the process^33^.

### *E. coli* CLC opens β_2_ by crab claw-like motions

In the CLC•β_2_^open^ structure, β_2_ is open with a gap of ∼20 Å, large enough for dsDNA to enter (Fig. 3a). This structure represents the stage where β_2_ is fully opened and ready to be transferred to p/t DNA (Fig. 1d, state **C**). The structure is similar to the yeast RFC•PCNA^open^ structures^33^ and the crystal structure of *E. coli* δγ_3_δ’ (without β_2_) bound to p/t DNA (CLC•DNA; PDB 3GLI)^27^. The CLC in the two *E. coli* structures, with and without bound β_2_, superimpose with an RMSD of 1.09 Å over 1417 Cα atoms across all five subunits. Compared to CLC•β_2_^closed^, the AAA+ modules of CLC•β_2_^open^ are more tightly packed (Figs. 2b and 3b). The total interface area between the AAA+ modules increases by 52% from 2781 Å^2^ in CLC·β_2_^closed^ to 4225 Å^2^. The most dramatic change occurs at the γ1γ2 interface with the buried area increasing by more than 900 Å^2^ (Fig. 3b), transforming the most loosely packed interface to the most tightly packed. There is also a significant increase at the γ2γ3 interface, whereas the δγ1 interface barely changes and these two modules move almost as a rigid body. In the transition from CLC•β_2_^closed^ to CLC•β_2_^open^, the chamber of the CLC expands mainly by outward rotation of the AAA+ modules of δ and γ1 pivoting around that of γ2 (Fig. 3c).

The β_2_ ring opens mostly by in-plane outward rotation of individual β domains and remains mostly flat after opening. Significant movements occur between Domains I and II of the β-I subunit that move outwards by 14°, and at the unbroken β_2_ interface that swings out by 11°, together providing greatest leverage for clamp opening at the opposite interface. A smaller change occurs between Domains I and II of β-II (Fig. 3d). Domains II and III of both β subunits move mostly as rigid bodies. Their connection is either rigid or held together by δ and γ3 binding in the canonical CBM peptide-binding sites.

During opening of β_2_, the AAA+ module of γ3 holds onto Domains II and III of β-II, while that of δ bound tightly to Domains II and III of β-I rotates outwards, causing rotation of β-I that pulls it away from β-II (Fig. 3c, d). Upon breaking of the dimer interface, Domain I of β-II also extends to release spring tension of the closed clamp^15^, leading to wider opening. During the opening, all CLC subunits interact with β_2_ more and more extensively until they tightly hold the clamp open. 3DVA of all “open” particles for the last 3D classification also captured similar movements at the final stage of clamp opening from partly to fully open (Fig. 3e). Therefore, the *E. coli* CLC opens β_2_ by a “crab-claw” mechanism.

### All CLC core subunits bind to the clamp in a similar fashion

Although CLCβ_2_ interactions are important for clamp loading and all of δ, γ and δ’ can interact with β^44^, to date only the δβ interaction has been visualised^15^. In CLC•β_2_^open^, all five subunits interact extensively with β_2_ (Fig. 4a, b), burying interface areas of 757, 709, 382, 989 and 847 Å^2^ between β and δ, γ1, γ2, γ3 and δ’, respectively (Extended Data Table 3). Like δ, all CLC subunits bind to surface crevices between pairs of structurally similar β domains through peptides located at the C-terminal ends of clamp-interacting helices and the following loops. The δ subunit binds to the canonical peptide-binding site on β-I *via* peptide 65-FSLC<u>QAMSLF</u> similar to that seen in CLC•β_2_^closed^ and the structure of δβ^15^. The γ1 subunit binds to a site between Domains I and II of β-I and γ2 binds mainly to Domain I of β-I near the dimer interface, while γ3 binds to the second CBM-binding site on β-II. Binding of the γ subunits to β_2_ is primarily mediated by different parts of peptides 109-DNVQYAPAR. In each case, residue Tyr113 of γ serves as an anchor for binding by nestling in a cavity in β_2_. Peptide 86-RFVLIEI of γ3 provides additional nearby contacts with β, and δ’ binds between domains I and II of β-II through peptide 66-QLMQAGTHP at a site involving the outside rim surface of β_2_ and peptide 101-NEHARLG at a nearby site. In comparison, the peptide of δ fits more snugly in its binding site than others and the δβ interface is more hydrophobic, contributing more binding energy than the other interfaces, which are more polar and transient (Extended Data Table 3).

**Fig. 4.**
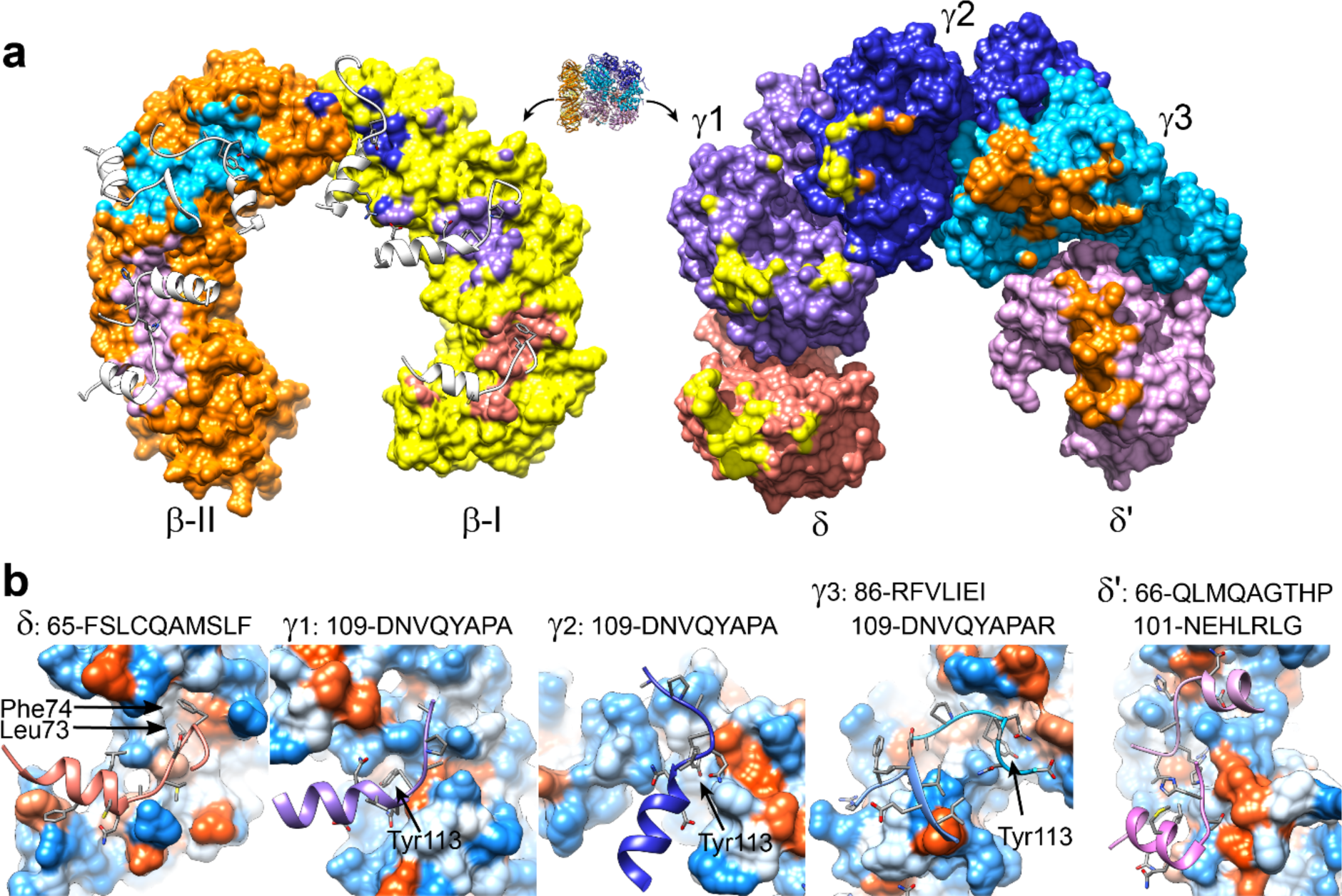
All CLC core subunits bind to domains of the open clamp in a similar fashion. **a**, CLCβ interfaces on β_2_ and the CLC of CLC•β_2_^open^. CLCβ interactions are mediated by peptides of the CLC subunits that are presented on the surfaces of β_2_ with their secondary structures depicted (white). Interface areas are coloured as their interacting counterparts. Peptide-binding surfaces of the δ, γ3 and δ’ subunits on β_2_ are more extensive than those of γ1 and γ2. **b**, Close-up views of clamp-binding peptides in binding pockets of β_2_. Sequences of peptides involving in clamp binding are shown above. Important residues, Leu73 and Phe74 of δ and Tyr113 of γ subunits that anchor the peptides in the binding sites, are labelled. The surface of β_2_ is coloured by hydrophobicity with hydrophobic areas in orange, neutral areas in white and hydrophilic areas in blue.

### Closing of β_2_ around a p/t DNA junction

To delineate the events after p/t DNA binding, we determine structures of CLC·β_2_ on p/t DNAs. First, we tested how different nucleotides affected the stability of the CLC•p/t DNA complex by SPR. We found that the CLC did not bind stably in the presence of ATP to a p/t DNA containing 23 bp dsDNA and a 5’-dT_50_ overhang (to which single-stranded DNA-binding protein SSB can also bind) (Extended Data Fig. 3a), as expected since p/t DNA-stimulated ATP hydrolysis by the CLC helps to quickly eject it from the DNA^19,29–32,39,45^. While the CLC with either ATPγS or ADP•AlF_x_ bound to p/t DNA, the ADP•AlF_x_- bound CLC•p/t DNA complex was very much more stable than that with ATPγS (Extended Data Fig. 3b−f). This suggests that ADP•AlF_x_ may acts truly as an analogue of the transition state in ATP hydrolysis^40^, where it stabilises the ATP-binding sites and freezes the whole CLC•p/t DNA complex. Moreover, both the β_2_ clamp and SSB were able to bind sequentially to the very stable ADP•AlF_x_-bound CLC•p/t DNA complex, forming very stable complexes with both the γ_3_ and τ_3_ CLCs (Extended Data Fig. 3g, h).

Therefore, samples containing the CLC (δτ_3_δ’ψχ instead of the γ_3_ CLC used previously), β_2_, ADP•AlF_x_ and p/t DNAs containing a 23 bp dsDNA and either a 5’-dT_10_ overhang or 5’-dT_45_ overhang plus SSB (that can bind more stably to dT_45_ than to dT_10_) were used for cryo-EM studies. In addition, a sample of the δτ_3_δ’ψχ CLC alone with ADP•AlF_x_ and a p/t DNA containing 15 bp of dsDNA and 5’-dT_10_ overhang was used. These complexes were stable and did not require crosslinking for data collection. Note that although the τ_3_-CLC was used for all following complexes with p/t DNA, in no case did we observe any additional cryo-EM density beyond that for the common Domains I–III of γ/τ seen in the γ_3_-CLC structures above. For consistency, these regions are referred to hereafter as γ rather than τ.

While samples with the 5’-dT_10_ overhang and no SSB contained only CLC bound to a closed β_2_ clamp (CLC•DNA·β_2_^closed^; solved at 2.6 Å resolution; Extended Data Fig. 4), samples with 5’-dT_45_ and SSB contained CLCs bound either to a partially open (CLC•DNA·β_2_^open^) or a closed (CLC•DNA·β_2_^closed^) clamp (Fig. 5a, b, Extended Data Table 1 and Fig. 5). The CLC•DNA·β_2_^open^ dataset was further separated into two similar complexes by 3D classification with β_2_ opened by ∼8 Å (CLC•DNA·β_2_^open^_1_; 2.9 Å resolution) and ∼6 Å (CLC•DNA•β_2_^open^_2_; 3.0 Å). These two structures are similar, representing equilibrated intermediates in the transition-state structure that occur prior to complete closure of β_2_ around DNA. Only CLC•DNA·β_2_^open^_2_ will be discussed.

**Fig. 5.**
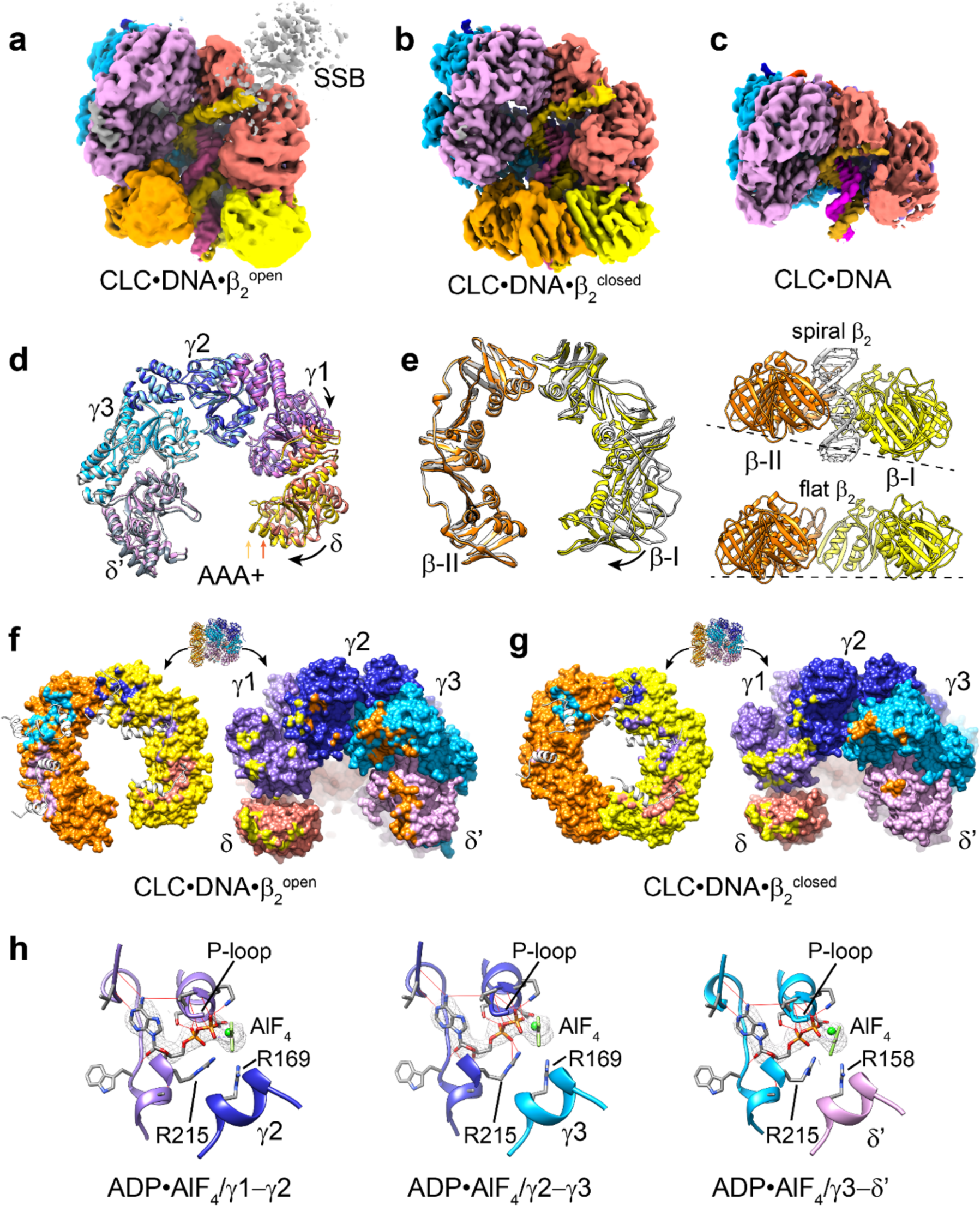
The β_2_ clamp partially or completely closes on loading onto primer-template DNA. **a**, Cryo-EM map of the CLC bound to a partially open clamp and p/t DNA (CLC•DNA•β_2_^open^). Densities corresponding to individual subunits are coloured as in Fig. 2. Density of the template DNA strand is coloured gold and the primer stand pink. Grey density at the top right corner belongs to SSB. **b**, Cryo-EM map of the CLC bound to a closed clamp and p/t DNA (CLC•DNA•β_2_^closed^). **c**, Cryo-EM map of the CLC bound to p/t DNA (CLC•DNA). **d**, The CLC constricts slightly on binding of p/t DNA. The AAA+ modules of CLC•β_2_^open^ and CLC•DNA•β_2_^open^ (coloured differently) are superimposed to show small inward movements of the CLC upon p/t DNA binding. Arrows point to the tips of the δ subunits. **e**, Orthogonal views showing β_2_ partially closes on p/t DNA and also becomes spiral. β_2_ of CLC•β_2_^open^ in the left pane is coloured grey and CLC•DNA•β_2_^open^ yellow and orange. Most movement occurs in the β-I subunit. β_2_ of CLC•DNA•β_2_^open^ becomes lock washer-like (right). **f**, **g**, CLCβ interfaces on β_2_ and CLC of CLC•DNA•β_2_^open^ and CLC•DNA•β_2_^closed^, respectively, showing that the latter has much reduced contacts. **h**, Mg•ADP•AlF_4−_ molecules are present in all three ATP-binding sites of CLC•DNA•β_2_^closed^ and the arginine finger residues: R169 of γ2 and γ3 and R158 of δ’, coordinate the AlF_4−_ groups that mimic the γ-phosphates of ATP. Mg^2+^ (spheres) and AlF_4–_ ions are coloured in shades of green and EM densities of Mg•ADP•AlF_4−_ are shown as mesh. Hydrogen bonds between ADP•AlF ^−^ and residues of the CLC are shown as red lines.

The structure of the CLC on p/t DNA alone (CLC•DNA; 2.6 Å) (Fig. 5c and Extended Data Fig. 6) is similar to CLC•DNA•β_2_^open^, except density for Domain I of δ is more poorly resolved in the absence of the β_2_ interaction. In both structures, the entire dsDNA and six dTs of the 5’ overhang are resolved, longer than the four nucleotides seen in the crystal structure of the δγ_3_δ’ CLC•DNA^27^. The aromatic ring of Trp279 of δ that is conserved and important for p/t DNA binding^46^ sandwiches between the last two nucleotides of the 5’ ssDNA overhang that protrudes from the dsDNA-binding chamber of the CLC, *i.e.*, the fifth and sixth unpaired nucleotides from the p/t junction (dT^5th^ and dT^6th^) (Fig. 6a, b). The CLC mainly interacts with the template DNA strand as described for the earlier δγ_3_δ’ CLC•DNA structure^27^ and β_2_ has only limited contacts with DNA. The AlF_4−_ moiety of the ATP analogue ADP•AlF_4−_ is clearly visible in all three ATP binding sites in all structures (Fig. 5h). That none of the sites contain only ADP suggests the stabilised transition state conformations we observe are indicative of the concerted (rapidly sequential or simultaneous) hydrolysis of ATP at all three sites.

**Fig. 6.**
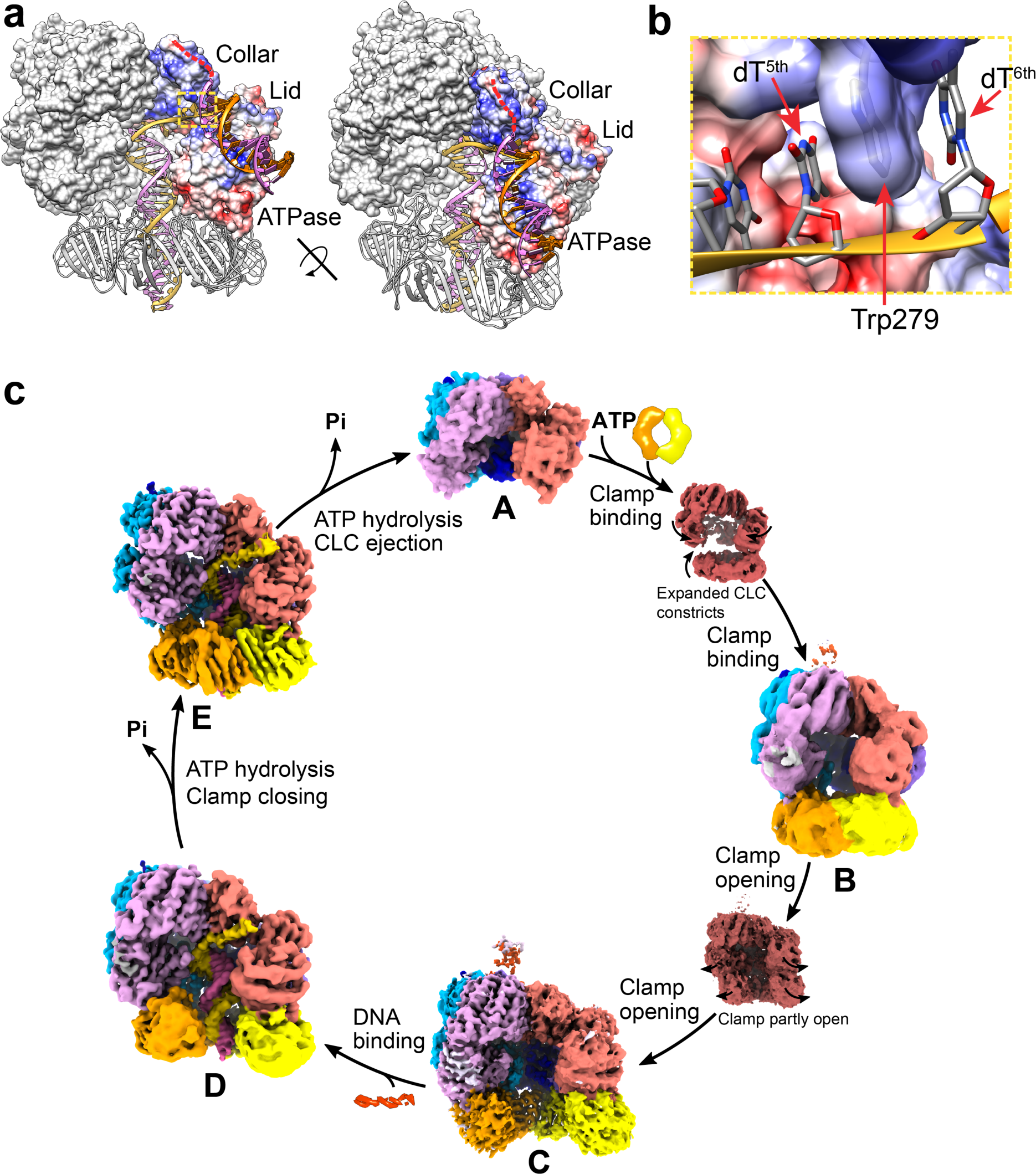
Structural views of clamp loading in *E. coli*. **a**, Recessed 5’-DNAs of nicked and gapped DNAs may bind to the exterior surface of δ. The exterior surfaces of the ATPase and collar domains of δ are mostly positively charged (blue) that suit DNA binding. If the template strand (orange) stays on the same path, the recessed 5’-DNA can bind to the ATPase domain and the unwound region (red dashed lines) can bind to the collar domain. The surface of δ is coloured by “Coulombic Surface Coloring” in Chimera^58,59^ that plots electrostatic potential. Positively charged areas are in blue, neutral areas white and negatively charged areas red. **b**, The aromatic side chain of Trp279 of δ that sandwiches between the bases of dT^5th^ and dT^6th^ of the ssDNA template may serve as a separation pin in unwinding the 5’-recessed ends of nicked and gapped DNAs to facilitate β_2_ loading. **c**, Schematic of the clamp loading pathway in *E. coli*. To load a clamp, the ATP-bound CLC (**A**) first binds to it *via* the δ subunit. The CLC then constricts, holds onto the clamp by simultaneous binding of the δ, γ1, γ3 and δ’ subunits (**B**). The CLC next expands and opens the clamp (**C**). The CLC−open clamp complex then directly binds to a p/t DNA. The CLC slightly constricts, and the clamp first partly closes (**D**) before complete closure around the p/t DNA junction (**E**). ATP hydrolysis, which may occur at any stage after p/t junction recognition, is presumed to accelerate clamp closing and trigger ejection of the CLC to allow a DNA polymerase to bind to the loaded clamp for DNA synthesis.

The CLCs in CLC•DNA, CLC•DNA·β_2_^open^ and CLC•DNA·β_2_^closed^ are very similar to one another and to the crystal structure of *E. coli* CLC•DNA^27^; they superimpose with CLC•DNA•β_2_^open^ with an RMSD of 1.13 Å over 1618 Cα atoms. Domain I of δ and the tips of β_2_ subunits in the CLC•DNA•β_2_^open^ structures are more poorly ordered, suggesting that either these parts are more flexible or the reconstructions contain a mixture of similar conformations. 3DVA of all refined particles with dT_45_ and SSB revealed that these reconstructions are indeed intermediates in an ensemble of continuously changing conformations. Compared to CLC•β_2_^open^, the CLC on DNA is slightly constricted (Fig. 5d). While transitioning from CLC•β_2_^open^ to CLC•DNA•β_2_^open^, the β-I subunit moves inwards and slightly downwards, so that β_2_ becomes spiral (Fig. 5e). For β_2_ to close around DNA, Domain I of δ moves farther inwards, allowing β-I to approach β-II, while β-II detaches from δ’, drops lower and re-closes the ring. The closed β_2_ is slightly distorted compared to the free clamp, with β-I more extended, as seen also in the structure of βδ^15^. While the CLC of CLC•DNA•β_2_^open^ maintains extensive interactions with β_2_ like in CLC•β_2_^open^, the interactions are much reduced in CLC•DNA•β_2_^closed^ (Fig. 5f and Extended Data Fig. 4e, f). The total interface area between the CLC and β_2_ decreases from 3663 Å^2^ in CLC•DNA•β_2_^open^ to 2496 Å^2^, which is nicely compensated by the reformation of a β dimer interface (667 Å^2^) and more contacts with DNA.

### Tethered SSB is distant from CLC and does not affect clamp closure on DNA

SSB can bind to the 5’ ssDNA overhang of a p/t DNA and interacts with χ and many other proteins *via* a conserved protein-interacting motif comprising the C-terminal 9 residues: 169-MDFDDDIPF^10,47^ (referred to as the C-tail, SSB-Ct). On p/t DNA with a 5’ overhang long enough to stably bind SSB (*e.g.*, 35 Nt or longer), SSB modestly stabilises the CLC on p/t DNA through interactions with χ, and binding of the CLC to a preformed p/t DNA-SSB complex leads to rapid remodelling of SSB to move it farther from the 3’ end of the primer, presumably to accommodate interaction of the δ subunit with the immediately-downstream ssDNA^48^. In all our structures with SSB, the presence of SSB is evident (Fig. 5a) but appeared to diffuse within a volume around the CLC at similar distances, as if being held remote from the CLC, perhaps by electrostatic repulsion. Surprisingly, there is no direct contact seen although SSB must be tethered to the CLC *via* interactions with χ^48–50^ and the ssDNA overhang.

We tried to uncover why β_2_ showed dynamic open and closed states after loading only in the presence of SSB and the dT_45_ overhang. To test if it is the interactions of SSB with χ or other CLC proteins, SSB binding to the 5’ ssDNA overhang, or the physical connection between the template DNA strand and ψχ mediated by SSB that enables β_2_ dynamics, several χ and SSB variants that are impaired in protein-protein interaction were used for cryo-EM studies. These proteins include χ∼SSB-Ct with the C-tail of SSB fused to χ by a flexible 9-residue linker that renders χ unable to interact with SSB^51^, SSB-T* containing only the ssDNA-binding structured OB-fold of SSB^52^, and SSB^CysApp^ with a Cys residue appended to the C-terminus of SSB that prevents protein interaction through SSB-Ct^53^. Five combinations in the CLC•DNA•β_2_ complex with SSB variants were tested: (i) ψχ∼SSB-Ct without SSB; (ii) ψχ∼SSB-Ct plus SSB; (iii) ψχ∼SSB-Ct plus SSB-T*; (iv) ψχ plus SSB-T*; and (v) ψχ plus SSB^cysApp^. All five samples contained CLC complexes with both partially open and closed β_2_ in similar ratios of particles. Therefore, interaction of χ with SSB does not appear to affect β_2_ dynamics after DNA binding. Finally, we found that the dT_45_ ssDNA overhang alone was sufficient to keep a fraction of the clamp open. It seems likely therefore that the energy barrier between CLC•DNA•β_2_^open^ and CLC•DNA•β_2_^closed^ is so small that just a longer 5’ ssDNA overhang is enough affect this equilibrium.

## Discussion

The structures of *E. coli* CLC•β_2_ complexes off or on a p/t DNA presented here are critical pieces of the clamp loading puzzle. Now every step of clamp loading in *E. coli*, from clamp binding, opening and loading, has a representative structure. These structures have provided abundant molecular and mechanistic details and insights into the clamp loading process.

### *E. coli* CLC actively opens β_2_ by crab claw-like motions in a direction opposite to yeast RFC

*E. coli* β_2_ is a very stable dimeric ring that rarely opens in solution^12,13,37,38^. Although an early fluorescence study showed that the *E. coli* CLC only underwent limited conformational changes during clamp opening and may passively “template” an open β_226_, it is now certain that ATP binding to the CLC actively opens bound β_2_ clamps^22,25^. Here, we provide direct evidence that the CLC indeed opens the clamp in a process that does not require ATP hydrolysis to produce a gap wide enough to allow direct passage of dsDNA using a “crab-claw” mechanism. 3D variability analysis (3DVA) of our reported structures suggests that δ is the first subunit of the CLC to bind to β_2_. Upon binding *via* δ, β_2_ and the CLC gradually move closer to each other, while the CLC constricts until it can hold onto β_2_ by interactions with the δ, γ1, γ3 and δ’ subunits (Fig. 4b). In contrast to yeast RFC that is constricted even prior to PCNA binding, the conformation of the free *E. coli* CLC appears more flexible and likely can spontaneously expand and constrict. Although δ binding alone is not sufficient to open β_2_, it must help to destabilise the clamp. Similar to the crystal structure of βδ^15^, the β-I subunit in CLC•DNA•β_2_^closed^ is more extended than in the free clamp, and this may indicate some spring tension in the closed β_2_ ring that distorts it and weakens the subunit interface. The CLC then undergoes further conformational changes in which its AAA+ domains become tightly packed, especially at the γ1γ2 interface. As a result, the AAA+ modules of δ and γ1 rotate outwards as a rigid body while γ3 holds the β-II subunit stationary, causing the β-I subunit to be pulled away in a hinge-like motion. Such a mechanism is consistent with the proposed role of δ as a “wrench”^16^ that is actively involved in clamp opening, but it is γ3 rather than δ’ that has the important role as a “stator”^16^ in holding the β-II subunit.

Like the *E. coli* CLC, yeast RFC also opens PCNA using a crab-claw mechanism^33^. However, RFC opens PCNA with larger scale conformational changes in a direction opposite to that of the *E. coli* CLC. The AAA+ domains of *E. coli* CLC•β_2_^closed^ and yeast RFC•PCNA^closed^ are overtwisted with the gaps in otherwise symmetrical CLCs invaded. However, it is the AAA+ modules of subunits D and E in yeast RFC that are most displaced from “ideal” positions rather than those of A (δ) and B (γ1) in the *E. coli* CLC. Consequently, yeast RFC opens a PCNA clamp by moving AAA+ modules C, D and E outwards in a clockwise direction as viewed from the collar, while *E. coli* CLC opens β_2_ by moving A (δ) and B (γ1) anticlockwise. Like the yeast complexes, the conformational changes from CLC•β_2_^closed^ to CLC•β_2_^open^ are accompanied by substantial increases in protein-protein interactions both among CLC subunits and between the CLC and β_2_; hence clamp opening is primarily driven by the free energy of protein interactions. The crab-claw mechanism used by *E. coli* CLC is not contradicted by the results of a fluorescence assay that suggest the CLC only undergoes limited conformational changes during clamp opening^26^. First, the conformational changes of the CLC are indeed smaller than yeast RFC. Secondly, the distances between the FRET pairs introduced in the CLC fall between those on CLC•β_2_^closed^ and CLC•β_2_^open^. To open β_2_, CLC first constricts to bind, then expands to open. The net distance changes between the fluorophores from free CLC to CLC•β_2_^closed^ then CLC•β_2_^open^ are too small to be accurately detected.

### *E. coli* CLC directly loads clamps onto p/t DNA junctions

A “filter and slide” mechanism of clamp loading onto p/t DNA was proposed based on the crystal structure of the T4 CLC•DNA•clamp^open^ complex^4,28^, in which only ssDNA can pass through the gap in the clamp and p/t DNA has to screw up the chamber of CLC for binding. However, it has since been shown that the *E. coli* CLC, yeast RFC and the Rad24-RFC^54^ that loads the 9-1-1 checkpoint clamp onto ssDNA or dsDNA with a 5’ recessed end all open their cognate clamps laterally with a gap large enough for dsDNA to enter. This may be a common characteristic in clamp opening, including in phage T4. Like the *E. coli* CLC•DNA•β_2_^open^ and yeast RFC•DNA•PCNA^open^ structures, the T4 CLC•DNA•clamp^open^ structure^4,28^ is likely a partially closed intermediate rather than a fully open complex. As the conformation of the *E. coli* CLC only changes slightly before (CLC•β_2_^open^) and after DNA binding (CLC•DNA•β_2_^open^) and the critical residue Tyr316 of δ that directly stacks on the 3’ base of the primer strand to recognise the p/t junction is well poised to interact, CLC•β_2_^open^ must be able to bind to a p/t junction directly. Compared to the “filter and slide” mechanism, direct clamp loading is more efficient and feasible since the *E. coli* CLC is tethered to the DnaB helicase through Domain IV of τ^10^ and hence is very close to priming sites at replication forks, perhaps not requiring an extensive search for a p/t terminus on ssDNA. The τ subunit can thus chaperone the CLC•β_2_^open^ complexes onto a newly primed site, then the Pol III core to the loaded clamp. Compared to the direct recognition of a p/t DNA junction by the *E. coli* CLC, yeast RFC has to partly melt the p/t junction, by which the 3’ base of the primer strand is flipped out to a pore where it stacks with Phe582 of Rfc1, a residue that is conserved in eukaryotes^33^. The p/t DNAs are also likely to bind to exterior sites on Rfc1 before being transferred into the central chamber of RFC, as do nicked and gapped DNAs^34–36^.

### How can the *E. coli* CLC bind to nicked and gapped DNA?

The *E. coli* CLC is also known to load β_2_ clamps onto nicked and gapped DNAs. However, the CLC can directly bind to p/t DNA and we have not yet observed dsDNA binding to sites other than in the central chamber. For yeast RFC, the 5’-recessed ends of nicked and gapped DNAs were found to bind to and partially melt on the collar domain and the N-terminal DNA-binding BRCT domain of Rfc1 that is flexibly attached to the main body of Rfc1^36^. This domain may help to search and bring DNA substrates to the RFC. It is imaginable that nicked and gapped DNAs may also bind to exterior sites on the *E. coli* CLC where the 5’-junctions are partly unwound. The flexible C-terminal region of τ is known to bind DNA^55^. In this study, we found that δτ_3_δ’ bound to p/t DNA much more quickly than δγ_3_δ’ (Extended Data Fig. 3i) and the unstructured region of γ binds back to δ’ near the entrance of the CLC (Fig. 3a). Therefore, the C-terminal region of τ may help to bind and bring in DNAs similarly to the BRCT domain of yeast Rfc1. The potential binding sites for the recessed 5’-DNA could be on the positively charged exterior surface of δ (Fig. 6a). In our structures, six nucleotides of the 5’ template overhang are seen to emerge from the CLC between the collar and ATPase domains of δ (Fig. 5b). This length of ssDNA is about optimal for gapped DNA to bind to yeast RFC^35^. Therefore, the outermost visible nucleotide, dT^6th^ that is the sixth unpaired nucleotide from the p/t junction, could either be at the 5’-ss/dsDNA junction or the last unpaired nucleotide of a nicked or gapped DNA. If the template strand stays on the path, the recessed 5’-DNA would bind to the ATPase domain of δ and the melted strand could bind to the collar domain (Fig. 6a). The aromatic ring of Trp279 of δ that is important for p/t DNA binding^46^ sandwiches between dT^5th^ and dT^6th^ (Fig. 6b), potentially serving as a separation pin to unwind and stabilise the 5’-junction.

### ATP hydrolysis in clamp closure around p/t junctions

It is well established that hydrolysis of the three bound molecules of ATP by the *E. coli* CLC occurs during clamp closure and subsequent ejection of the CLC^28–32^. In one of the crystal structures of T4 CLC•clamp^closed^ (PDB 3U61)^28^, the ATP-binding site at the A–B subunit interface is occupied by ADP. It was proposed that ATP hydrolysis at this site triggers conformational changes in the T4 CLC and clamp closure. In contrast, ATPγS was present in the four main ATP-binding sites in the yeast RFC•DNA•PCNA^closed^ complexes and all ATP-binding sites in an alternate T4 CLC•clamp^closed^ structure (PDB 3U5Z)^28^ are occupied by ADP•BeF_4_.

We expected that one or more of the ATP-binding sites in our CLC•DNA•β_2_^closed^ structure would be occupied by ADP while the other(s) are bound to the ATP analogue ADP•AlF_4−_, reflecting the order of ATP hydrolysis at the three sites during clamp closing. This was not the case. Indeed, in all of our structures of the CLC bound to p/t DNA (with or without β_2_), the ATP binding sites are fully occupied by ADP•AlF_4−_, consistent with this nucleotide acting as an analogue of the transition state on the pathway of ATP hydrolysis at all three sites (Fig. 5h; Extended Data Table 2). This observation suggests that ATP hydrolysis at all three sites occurs simultaneously (or nearly so) during clamp closing and CLC ejection. Moreover, rather than the trapping of the ATPase sites in transition state structures leading to a single intermediate state of the CLC and bound clamp, it leads to a delicately poised dynamic equilibrium between open, closed and intermediate state(s) of the CLC and the clamp itself.

### Distinct features of clamp loading in *E. coli*

Compared to eukaryotic RFCs, the *E. coli* CLC lacks an A’ domain and an E-plug that block dsDNA from entering the chamber prematurely^21,33^. It has been believed that the free *E. coli* CLC is disorganised in such a way not to suit high-affinity DNA binding^39^. We confirmed earlier studies that showed the ATP-bound *E. coli* CLC was unable to bind stably to p/t DNA^56^, while it formed very stable complexes with p/t DNA in the presence of ATPγS^27,56^ or ADP•AlF_x_ (Extended Data Fig. 3a–c). It seems most likely that the ATP-bound CLC appears to interact only transiently with p/t DNA in the absence of the clamp because DNA-stimulated ATP hydrolysis^19,39,45^ quickly ejects it. Probably, CLCs are bound to clamps most of the time in cells, or it is only through clamp binding and the “crab-claw” motions *en route* to β_2_ opening that the CLC becomes properly organised and ready to accommodate a p/t DNA. Secondly, bacterial clamps are homodimers; therefore, the CLC only has to stabilise one of the two clamp interfaces during clamp opening at the other. Thus, the *E. coli* CLC opens β_2_ by holding one subunit while rotating the other away. This keeps one interface closed while providing sufficient leverage to open the opposite interface. Upon opening, Domain I of β-II also becomes more flexible, facilitating dsDNA passage. In contrast, eukaryotic and phage CLCs have to keep two interfaces of their trimeric clamp stable with the assistance of an extra A’ domain^27,33^. Thus, dimeric bacterial clamps possess different CLC binding pockets located in each of the three domains of each subunit (Fig. 4), while trimeric clamps contain only alternate strong and weak sites. The *E. coli* CLC relies mostly on the two canonical peptide-binding sites in β_2_, where it binds subunits δ and γ3. These two sites are more hydrophobic and bind the CLC with higher surface complementarity (Extended Data Table 3). The γ subunits adapt to having three different types of binding sites by using different parts of the same peptide segment in each (Fig. 4b). As for interactions with DNA, while yeast RFC interacts with both template and primer strands^33^ and different eukaryotic clamp loaders require different A subunits for different DNA substrates^4,54^, the *E. coli* CLC mainly interacts with the template strand and is more promiscuous in that it can load clamps onto different DNA substrates for other essential activities, such as DNA repair. Despite these differences, the processes of clamp loading are very similar in *E. coli* and yeast and the general mechanisms are likely conserved in all life.

Based on these structures, we have delineated every step of the entire pathway of clamp loading in *E. coli* (Fig. 6c). We show that the ATP-bound *E. coli* CLC (**A**) first binds to the β_2_ clamp *via* the δ subunit. The CLC then constricts and interacts with β_2_ *via* simultaneous binding of the other CLC subunits (**B**). Next, the CLC expands and the AAA+ modules of the δ and γ1 subunits rotate outwards, thereby opening the β_2_ ring (**C**). The resulting CLC·β_2_^open^ complex directly binds or is transferred to the p/t DNA junction, in a process that may involve the C-terminal regions of τ. Next, the CLC slightly constricts to allow the clamp to first partly (**D**) then completely close around the p/t DNA junction (**E**). Finally, the CLC dissociates (**A**), leaving β_2_ at the p/t junction for the Pol III polymerase core to bind and DNA replication to start.

### Online content

Additional methods and references, Nature Research reporting summaries, source data, extended data, supplementary information, acknowledgements, peer review information; details of author contributions and competing interests; and statements of data and code availability are available at http://doi.org/…

## Methods

### Proteins and oligonucleotides

Proteins and complexes were prepared as previously reported: the δτ_3_δ’ and δγ_3_δ’^37^ minimal clamp loader complexes, ^bio^χψδτ_3_δ’ full clamp loader complex with N-terminally biotinylated χ^60^, the ψχ clamp loader accessory complex^61^, β_2_ sliding clamp^41^, single-stranded DNA-binding protein SSB^62^ and SSB-T*, its OB-domain^52^. Proteins χ∼SSB-Ct with the C-tail of SSB (MDFDDDIPF) fused to χ by a flexible 9-residue linker (TRESGSIGS)^51^, and SSB^CysApp^ with a cysteine residue appended to the C-terminus of SSB^53^ were isolated similarly to the wild-type proteins^60,62^.

Primer-template (p/t) DNAs (all from Integrated DNA Technologies, HPLC purified) for cryo-EM were prepared by annealing a primer oligonucleotide 862 (5’-GAGATAGTTACAACATACGATCG) to template strands 864 (5’-T_10_-CGATCGTATGTTGTAACTATCTC, for experiments of CLC•DNA•β_2_ without SSB) or 863 (5’-T_45_-CGATCGTATGTTGTAACTATCTC; for experiments of CLC•DNA•β_2_ with SSB); and 629 (5’-TTAGTTACAACATACT) to 883 (5’-T_10_-AGTATGTTGTAACTA) for of CLC•DNA. For SPR studies, the biotinylated template strand 865 (5’-biotin-dT_50_-CGATCGTATGTTGTAACTATCTC) was annealed to primer strand 862.

### Surface plasmon resonance

SPR experiments were carried out on a Biacore T200 instrument (Cytiva) operated at 20°C using a streptavidin-coated (SA) chip (Cytiva). The SPR buffer contained 30 mM Tris-HCl pH 7.6, 400 μM ATP, 8 mM MgCl_2_, 30 mM NaCl, 0.5 mM dithiothreitol, 0.005% surfactant P20; or 30 mM Bis-Tris pH 6.4, 0.2 mM ADP, 3 mM NaF, 0.3 mM AlCl_3_, 8 mM MgCl_2_, 30 mM NaCl, 0.5 mM dithiothreitol, 0.005% surfactant P20. Bis-Tris at pH 6.4 was used to increase the solubility of AlF_x_.

For interaction of β_2_ with the immobilized CLC complex, ^bio^χψδτ_3_δ’ or ^bio^χψδγ_3_δ’ containing N-terminally biotinylated χ (20 nM) was loaded onto a flow-cell of the SA chip to achieve a response of ∼1400 RU. Serially diluted solutions of β_2_ in buffer containing desired nucleotides were made to flow over the surface at 10 μl/min for desired durations. The bound protein was allowed to dissociate until the signal nearly returned to baseline. To determine the *K*_D_ value of ^bio^χψδτ_3_δ’β_2_ interaction by kinetics fitting (Extended Data Fig. 1a), β_2_ was injected for 40 s, then its dissociation from the immobilized CLC was monitored for 700 s. The entire set of zero-subtracted data were fit simultaneously (globally) using a 1:1 Langmuir binding model to derive association and dissociation rate constants, *k*_a_ and *k*_d_. The dissociation constant *K*_D_ was calculated from Equation 1:

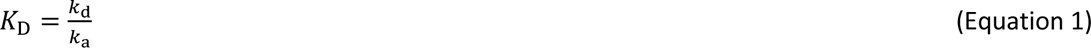

To determine *K*_D_ values of the ^bio^χψδγ_3_δ’β_2_ interaction by fitting using a 1:1 steady-state affinity (SSA) model (Extended Data Fig. 1b–d), β_2_ was injected for durations required to reach equilibrium with ATP, 100 s with ATPγS (400 μM) or 40 s with ADP•AlF_x_. The equilibrium responses of β_2_ interacting with the immobilized ^bio^χψδγ_3_δ’ were fit using using Equation 2:

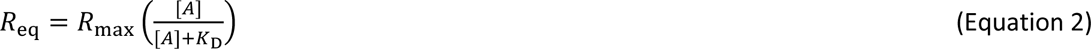

where R_max_ corresponds to the response when all immobilised ligand on the surface (^bio^χψδγ_3_δ’) is saturated with the analyte (β_2_), *K*_D_ is the disassociation constant and [*A*] is the concentration of analyte in solution.

For interaction of χψδγ_3_δ’ with immobilized p/t DNA (Extended Data Fig. 3), the biotinylated oligo 865 (3 nM) was loaded onto flow-cell 2 for 108 s, yielding a response of 85 RU. Oligo 862 at 1 μM was then injected to assemble p/t DNA on the chip surface *in situ*. This DNA template is comprised of a 23-bp dsDNA region positioned away from the surface and a (dT)_50_ 5’ ssDNA overhang attached to the chip surface through the stable biotin-streptavidin interaction. CLC (δγ_3_δ’ψχ, 200 nM) in buffer containing different nucleotides was injected for desired durations and the bound proteins were allowed to dissociate in buffer containing desired nucleotides. For assembly of the CLC complex on p/t DNA, 100 nM δγ_3_δ’ or δτ_3_δ’, 200 nM ψχ, 1.6 μM β_2_ and 20 nM SSB were injected sequentially.

### Sample preparation for cryo-EM

To prepare samples for cryo-EM, 30 μL of 6 μM δγ_3_δ’ or δτ_3_δ’ was mixed with ψχ complex at a molar ratio of 1:1.2, β_2_ at 1:1.3, p/t DNA (annealed DNA oligonucleotides) at 1:1.3 if required, and dialysed twice at 4°C against 250 mL of 30 mM Na.HEPES pH 7.5, 3 mM MgCl_2_, 2 mM dithiothreitol, 0.25 mM EDTA, 2% glycerol. Nucleotides (1 mM ATPγS or 2 mM ADP plus 5 mM NaF and 0.5 mM AlCl_3_) were then added to the dialysates. Samples without DNA were crosslinked by treatment with 2 or 5 mM bis(sulfosuccinimidyl)suberate (BS^3^) on the bench (∼21°C) for 15 min. Reactions were quenched by adding 1 M Tris-HCl pH 7.6 to 25 mM. Samples (3 μL) were applied onto Quantifoil UltrAuFoil grids (R1.2/1.3) that were blotted for 4 s at 6°C without force, then plunged into liquid ethane using a FEI Vitrobot Mark IV.

### Data collection, image processing and model building

Micrographs were collected using a Thermo Fisher Titan Krios G3i microscope at 300 kV, equipped with a Gatan K2 camera in electron counting mode at a nominal defocus range of −0.4 to −1.3 μm, with a pixel size of 0.82 Å/pixel and a total dose of 50 e^−^ over 5 s fractionated across 50 frames, except that a pixel size of 0.66 Å/pixel and a total dose of 60 e^−^ over 6 s fractionated across 60 frames was used for the sample containing 23-bp dsDNA and a dT_10_ 5’-overhang for the reconstruction of CLC•DNA•β_2_^closed^. Images were collected without use of an energy filter.

For samples without DNA, movies were corrected for drift and aligned using MOTIONCOR2^63^ with electron-dose weighting in RELION 3.1^64^. Motion-corrected micrographs were then imported into cryoSPARC^65^. CTF parameters were estimated with CTFFIND4.1^66^ and micrographs with estimated resolutions worse than 3.5 Å were excluded. Particles were first picked by blob-picker, extracted at 256-pixel box-size and subjected to 2D classification; classes showing high-resolution features were retained and used for Topaz^67^ training. The best Topaz training model was used to pick particles on all the micrographs. Particles were subjected to a round of 2D classification and good particles were retained for 3D classification by a combination of *ab-initio* reconstruction and heterogeneous refinement. Particles of selected 3D classes were used for homogeneous 3D refinement. Refined particles were imported into RELION 3.1, extracted at 360-pixel box-size for 3D auto-refinement, CTF refinement, Bayesian polishing and final 3D auto-refinement and postprocess. For samples with DNA, data were processed in RELION 3.1. Particles were first picked from a subset of micrographs by Laplacian picking and subjected to 2D classification. Selected 2D classes were then used for template picking on all micrographs. Particles thus picked were classified by rounds of Class2D and Class3D. Particles from selected classes were used for 3D auto-refinement, followed by CTF refinement, Bayesian polishing, then final 3D auto-refinement and postprocess. Particles of the final 3D reconstructions were imported into cryoSPARC for 3D variance analysis (3DVA). Global resolution is reported according to the 0.143 FSC criterion. Local resolutions of models were estimated using the RELION algorithm.

For model building, the structures of *E. coli* δγ_3_δ’•DNA complex^27^ and β_241_ were placed into density maps using Phenix.dock_in_map^68^. The resulting complexes were then refined in Phenix.real_space_refine^68^. After inspection and model building in COOT^69^, ATPγS or ADP•AlF_4−_ was included in the models. The models were subjected to rebuilding in COOT and REFMAC^70^ refinement, followed by a final cycle of refinement in Phenix.real_space_refine.

Molecular graphics and analyses used Chimera^58^ and ChimeraX^59^. Protein interface areas were calculated by the PDBePisa server^71^.

## Data availability

Cryo-EM maps and atomic coordinates have been deposited with the Electron Microscopy Data Bank and Protein Data Bank under accession codes EMD-40079 and 8GIY for CLC•β_2_^closed^, EMD-40080 and 8GIZ for CLC•β_2_^open^, EMD-40081 and 8GJ0 for CLC•DNA•β_2_^open^_1_, EMD-40082 and 8GJ1 for CLC•DNA•β_2_^open^_2_, EMD-40083 and 8GJ2 for CLC•DNA•β_2_^closed^, and EMD-40084 and 8GJ3 for CLC•DNA.

## Acknowledgements

We thank Yao Wang and Amy McGrath for gifts of proteins, and Jacob Lewis and Sarah Henrikus for discussion on cryo-EM data processing. We are especially grateful for mechanistic discussions with Linda Bloom. This work was supported by the Australian Research Council (DP180100805 to A. J. O. and N. E. D.; DP210100365 to H.G., P. J. L., G. T., A. J. O. and N. E. D.).

## Author contributions

Conceptualization, Methodology, Z.-Q. X., S. J., H. G., P. J. L., G. T., A. J. O., N. E. D.; Investigation: Z.-Q. X. (EM), A. C. P., S. H. J. B., J. C. B., G. T. (EM support), A. J. O. (model building), Z.-Q. X. (structure analysis), S. J., Z.-Q. X., A. C. P. (SPR); Resources, S. J., Z.-Q. X., A. T. Y. L; Writing – Original Draft, Visualisation, Z.-Q. X., Writing – Review & Editing, Z.-Q. X., S. J., A. J. O., N. E. D.; All authors have read and approved the final version.

---

**Extended Data Table 1.** Details of cryo-EM data collection, reconstruction and model refinement.

| | CLC•$\beta_2^{\text{closed}}$ | CLC•$\beta_2^{\text{open}}$ | CLC•DNA•$\beta_2^{\text{open}}$ | | CLC•DNA•$\beta_2^{\text{close}}$ | CLC•DNA |
| --- | --- | --- | --- | --- | --- | --- |
|  |  |  |  |  | <sup>d</sup> |  |
| **Data collection and processing** |  |  |  |  |  |  |
| microscope | Titan Krios | Titan Krios | Titan Krios |  | Titan Krios | Titan Krios |
| voltage (kV) | 300 | 300 | 300 |  | 300 | 300 |
| electron dose ($\text{e}^-/\text{\AA}^2$) | 50 | 50 | 50 | | 60 | 50 |
| dose rate ($\text{e}^-/\text{\AA}^2/\text{fraction}$) | 1 | 1 | 1 | | 1 | 1 |
| detector | K2 | K2 | K2 |  | K2 | K2 |
| defocus range ($\mu\text{m}$) | 0.4–1.3 | 0.4–1.3 | 0.4–1.3 | | 0.4–1.3 | 0.4–1.3 |
| calibrated pixel size ($\text{\AA}$) | 0.82 | 0.82 | 0.84 | | 0.66 | 0.84 |
| micrographs (No.) | 7269 |  | 4226 |  | 1703 | 2909 |
| total extracted particles | 1376k |  | 1433k |  | 600k | 645k |
| final particles | 373k | 280k | 75k | 80k | 452k | 308k |
|  |  |  | Open1 | Open2 |  |  |
| symmetry imposed | C1 | C1 | C1 | C1 | C1 | C1 |
| resolution ($\text{\AA}$) | 3.7 | 2.7 | 2.9 | 3.0 | 2.6 | 2.8 |
| **Refinement** |  |  |  |  |  |  |
| RMS deviations |  |  |  |  |  |  |
| – bond lengths ($\text{\AA}$) | 0.002 | 0.002 | 0.002 | 0.002 | 0.002 | 0.002 |
| – bond angles ($^\circ$) | 0.420 | 0.459 | 0.507 | 0.520 | 0.503 | 0.542 |
| Ramachandran (%) |  |  |  |  |  |  |
| – outliers | 0.00 | 0.04 | 0.04 | 0.16 | 0.04 | 0.06 |
| – allowed | 1.71 | 3.09 | 1.95 | 2.10 | 1.63 | 2.02 |
| – favoured | 98.29 | 96.87 | 98.01 | 97.74 | 98.33 | 97.93 |
| rotamer outlier (%) |  |  | 1.37 | 2.40 | 0.57 | 1.48 |
| Molprobity score | 1.08 | 1.52 | 1.13 | 1.37 | 0.96 | 1.18 |
| Clashscore | 2.93 | 5.78 | 2.41 | 2.40 | 1.95 | 2.47 |

**Extended Data Table 2.** Interatomic distances probe the integrity of ATP-binding sites in the CLC complex.

| | CLC•$\beta_2^{\text{closed}}$ | CLC•$\beta_2^{\text{open}}$ | CLC•DNA•$\beta_2^{\text{open}}$ | | CLC•DNA•$\beta_2^{\text{closed}}$ | CLC•DNA |
| --- | --- | --- | --- | --- | --- | --- |
|  |  |  | Open1 | Open2 |  |  |
| **Map resolution (Å)** | 3.7 | 2.7 | 2.9 | 3.0 | 2.6 | 2.8 |
| **Distances (Å)\*** |  |  |  |  |  |  |
| $\gamma 1\text{-R215-C}\zeta - \gamma 1\text{-ATP-P}\gamma^*$ | 4.6 | 4.6 | 6.0 | 4.2 | 4.6 | 4.8 |
| $\gamma 2\text{-R169-C}\zeta - \gamma 1\text{-ATP-P}\gamma^*$ | 13.1 | 4.9 | 4.8 | 4.4 | 4.7 | 4.6 |
| $\gamma 2\text{-R215-C}\zeta - \gamma 2\text{-ATP-P}\gamma^*$ | 4.7 | 4.2 | 4.8 | 4.6 | 4.7 | 5.0 |
| $\gamma 3\text{-R169-C}\zeta - \gamma 2\text{-ATP-P}\gamma^*$ | 9.0 | 4.9 | 4.5 | 5.2 | 4.6 | 4.7 |
| $\gamma 3\text{-R215-C}\zeta - \gamma 3\text{-ATP-P}\gamma^*$ | 4.6 | 3.7 | 4.4 | 4.5 | 4.6 | 4.8 |
| $\delta'\text{-R158-C}\zeta - \gamma 3\text{-ATP-P}\gamma^*$ | 5.4 | 4.5 | 4.8 | 4.5 | 4.5 | 4.9 |

\*Distances are from the C $\zeta$ atoms of arginine finger ( $\gamma$ -R169 or $\delta'$ -R158) and sensor II ( $\gamma$ -R215) arginines and either the P $\gamma$ atom of ATPγS (structures without p/t DNA) or the Al atom of ADP•AlF $_4^-$ (structures with p/t DNA). Distances shown in red are indicative of well-formed ATP binding sites.

**Extended Data Table 3.** Interfaces between clamp loader and β subunits in the CLC•β_2_^open^ complex*.

| Chain1 | Chain 2 | Interface area ($\text{\AA}^2$) | $\Delta^iG$ (kcal/mol) | $\Delta^iG$ (*P*-value) | HB/SB |
| --- | --- | --- | --- | --- | --- |
| $\delta$ (A) | $\beta 1$ (H) | 756.8 | -7.9 | 0.250 | 5/1 |
| $\tau 1$ (B) | $\beta 1$ (H) | 708.9 | -3.0 | 0.584 | 9/8 |
| $\tau 2$ (C) | $\beta 1$ (H) | 287.4 | -3.1 | 0.329 | 2/0 |
| | $\beta 2$ (I) | 95.0 | -1.9 | 0.239 | 0/0 |
| $\tau 3$ (D) | $\beta 2$ (I) | 988.9 | -4.5 | 0.480 | 11/3 |
| $\delta'$ (E) | $\beta 2$ (I) | 847.4 | -0.1 | 0.692 | 7/1 |
| Total |  | 3684.4 |  |  |  |

**Extended Data Fig. 1.**
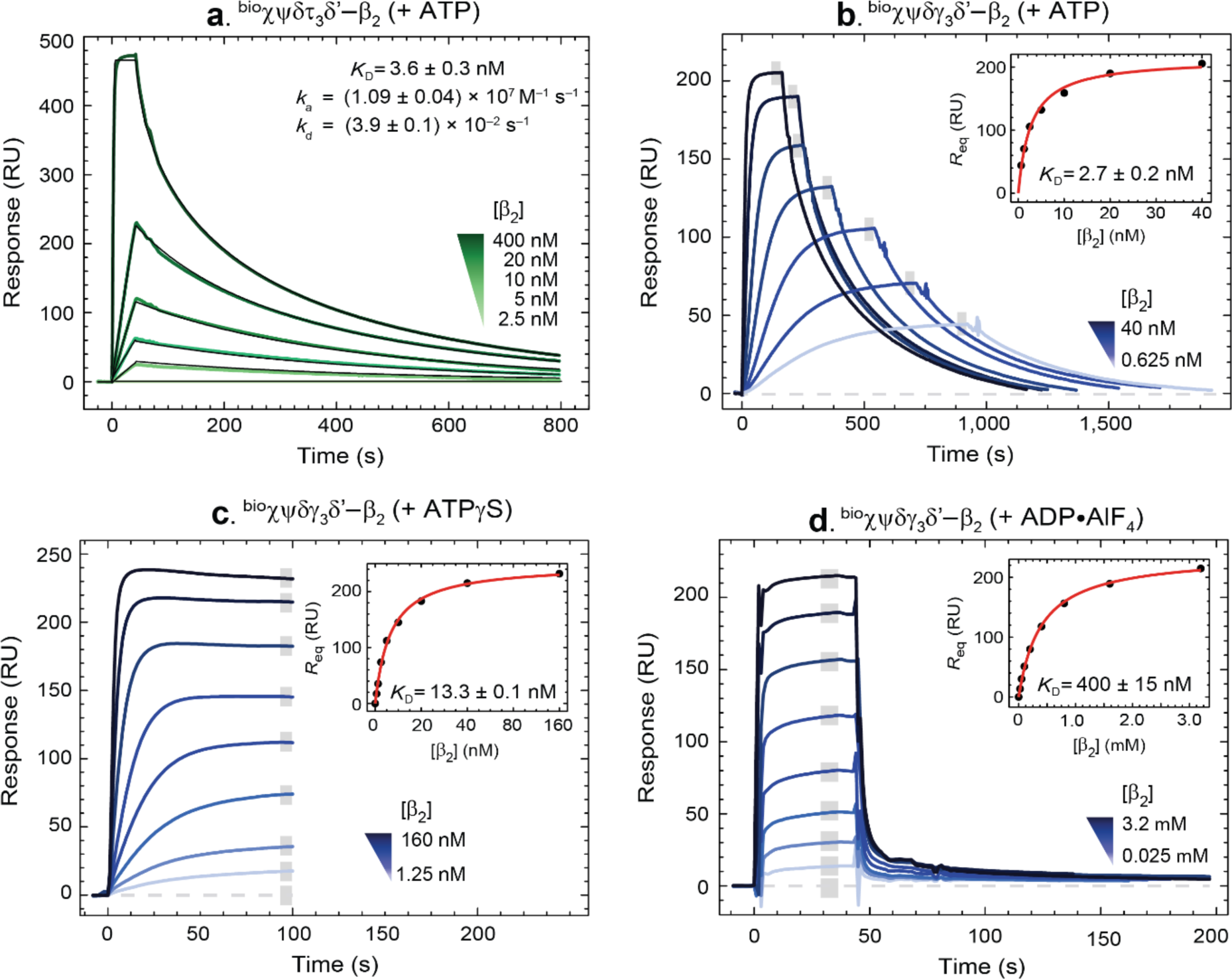
SPR study of interactions of the *E. coli* CLC with β_2_, related to Fig. 2. **a**, SPR sensorgrams show association (40 s) and dissociation (> 700 s) phases of β_2_ (2.5−20 nM and 400 nM) on ^bio^χψδτ_3_δ’ (including zero control) in the presence of 400 μM ATP. Curves (shown in shades of green), were fit simultaneously (black curves) to a Langmuir 1:1 binding model with mass transfer correction (LMT) to generate binding kinetics parameters: *k*_a_ and *k*_d_ (as shown); saturated binding level of β_2_, *R*_max_ = 476.8 ± 0.4 RU; and mass-transfer coefficient, *k*_t_ = (3.11 ± 0.01) x 10^8^ RU M^−1^s^−1^. *K*_D_ was then calculated from kinetics parameters using Equation 1. The high (diffusion controlled) value of *k*_a_ >> 1 x 10^6^ M^−1^s^−1^ for ternary ATP•^bio^χψδγ_3_δ’•β_2_ complex formation suggests significant mass transfer limitations, justifying use of the LMT model. **b**, SPR sensorgrams showing association and dissociation of β_2_ from ^bio^χψδγ_3_δ’ in the presence of 400 μM ATP over a 0.625−40 nM concentration range of serially diluted β_2_ samples. Sample solutions of β_2_ in SPR buffer were made to flow for durations sufficient to reach binding equilibrium (*R*_eq_). The responses at equilibrium, determined by averaging values in the grey bar region, were then fit (inset, red curves) using the steady-state affinity model (SSA) (Equation 2) to obtain a *K*_D_ value of 2.7 ± 0.2 nM. Note that the *K*_D_ values of ATP-dependent β_2_ binding to ^bio^χψδτ_3_δ’ (a) and ^bio^χψδγ_3_δ’ (b) are very similar, as expected. **c**, SPR sensorgrams showing the association phase (100 s) of β_2_ binding to ^bio^χψδγ_3_δ’ in the presence of 400 μM ATPγS over a 1.25−160 nM concentration range of serially diluted β_2_ samples (including 0 nM control). The responses at equilibrium, determined by averaging values in the grey bar region, were fit (inset, red curves) using the SSA model to obtain a *K*_D_ value of 13.3 ± 0.1 nM. **d**, SPR sensorgrams showing association (40 s) and dissociation of β_2_ from ^bio^χψδγ_3_δ’ in the presence of 400 μM ADP•AlF_x_ over a 0.025−3.2 mM concentration range of serially diluted β_2_ samples (including 0 mM control). The responses at equilibrium, determined by averaging values in the grey bar region, were fit (inset, red curves) using the SSA model to obtain a *K*_D_ value of 400 ± 15 nM. As it turns out, CLCβ_2_ interaction is much weaker with ADP•AlF_x_ compared to ATP or ATPγS. All errors are standard errors of the fits.

**Extended Data Fig. 2.**
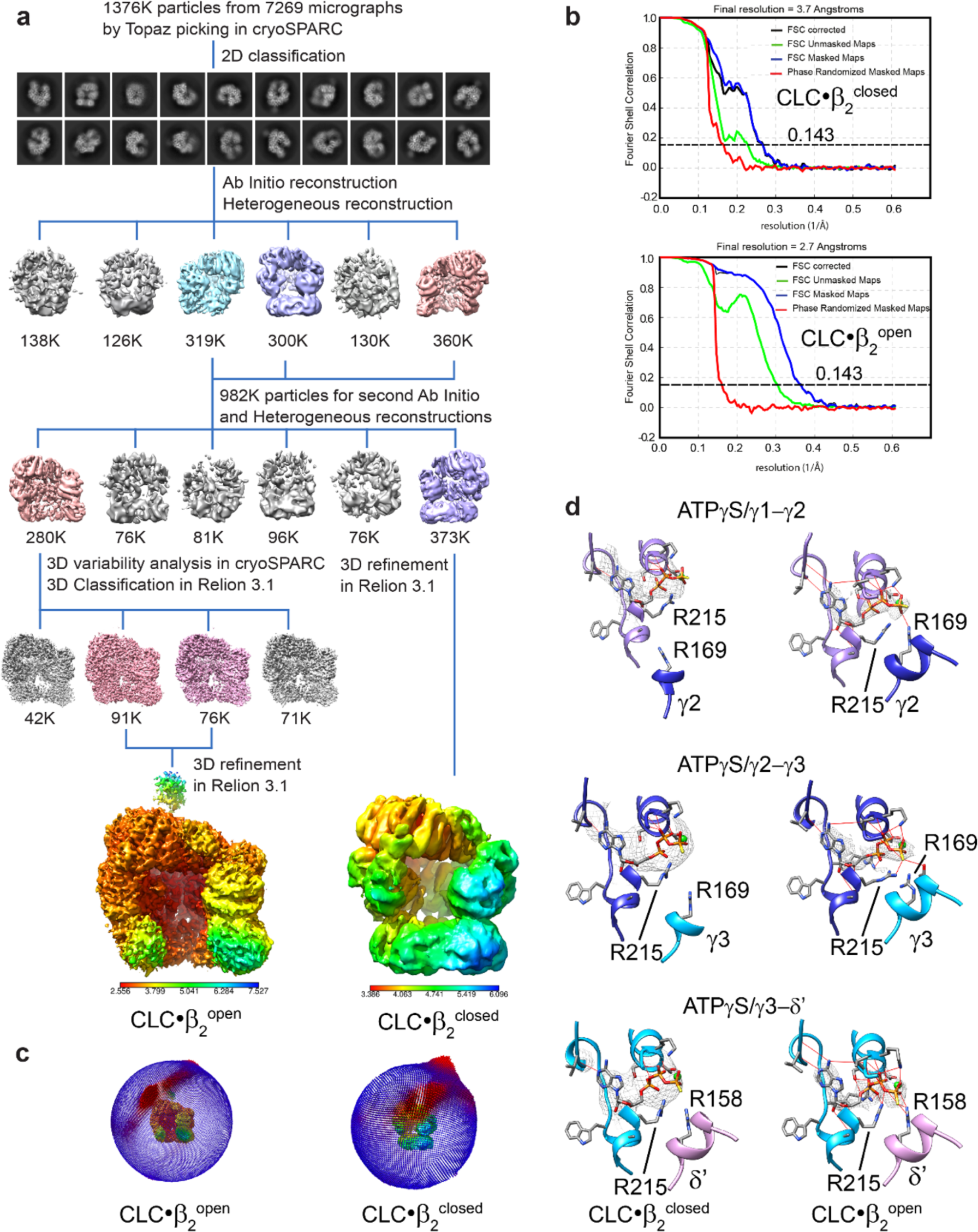
Processing of cryo-EM data of the CLC•β_2_^closed^ and CLC•β_2_^open^ complexes, related to Fig. 2, 3. **a**, Cryo-EM image processing workflow. Particle picking, 2D and 3D classifications, and initial 3D refinement were performed in cryoSPARC_65_. The refined particles of individual reconstructions were imported into RELION 3.1 for further 3D classifications, 3D refinement, CTF refinement and Bayesian polishing. **b**, Fourier shell correlation of the final density maps of CLC•β_2_^closed^ and CLC•β_2_^open^. **c**, Angular distribution plot of observed particle orientations in the reconstructions of CLC•β_2_^closed^ and CLC•β_2_^open^. **d**, Mg•ATPγS is present in all three ATP-binding sites of CLC•β_2_^closed^ and CLC•β_2_^closed^. Arginine fingers, Arg169 of γ2 and γ3 and Arg158 of δ’, are close to γ-phosphate groups of ATPγS in CLC•β_2_^closed^, but much farther away in CLC•β_2_^closed^. EM densities of ATPγS are shown as meshes. Hydrogen bonds between ATPγS and residues of the CLC are shown as red lines.

**Extended Data Fig. 3.**
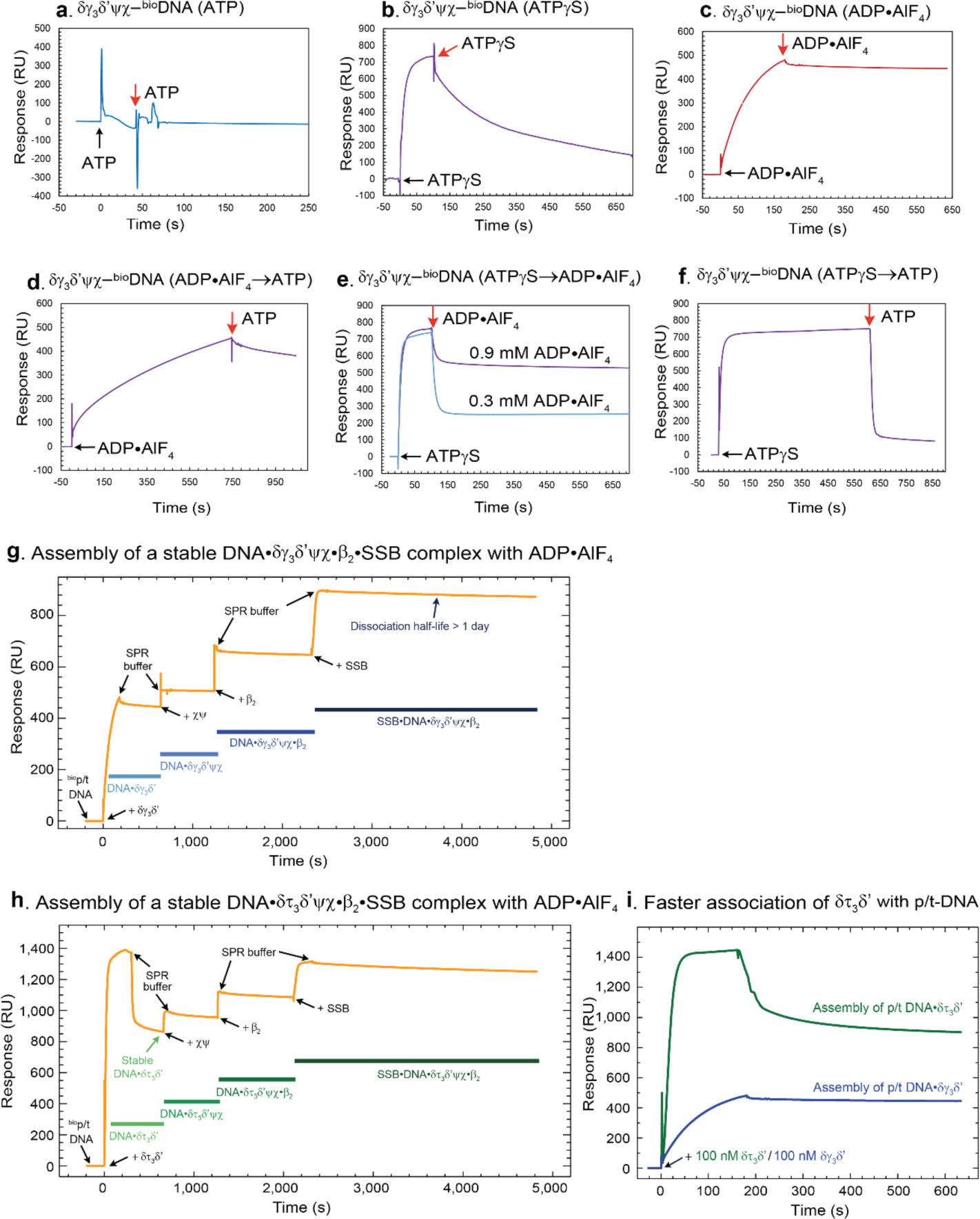
SPR study of interactions of the *E. coli* CLC with p/t DNA, related to Fig. 5. **a**−**c**, SPR sensorgrams showing the association and dissociation of δγ_3_δ’ψχ to and from p/t DNA that is immobilised on the surface of an SA chip *via* biotin in the presence of ATP, ATPγS and ADP•AlF_x_, respectively. Black arrows indicate the start of protein injections; red arrows indicate changes of buffers. **d**, SPR sensorgram of the association of δγ_3_δ’ψχ to p/t DNA with ADP•AlF_x_ and the dissociation with ATP, showing that ATP is slow to replace ADP•AlF_x_ from the CLC and the CLC•DNA complex remains quite stable. **e**, SPR sensorgram of the association of δγ_3_δ’ψχ to p/t DNA with ATPγS and the dissociation with 0.3 mM (blue) and 0.9 mM (pink) ADP•AlF_x_, respectively, showing that ADP•AlF_x_ replaced ATPγS from the CLC and the CLC•DNA complex became much more stable. **f**, SPR sensorgram of the association of δγ_3_δ’ψχ to p/t DNA with ATPγS and the dissociation with ATP, showing that ATP efficiently replaced ATPγS from the CLC and destabilised the CLC•DNA complex. **g**, Sensorgram showing responses as various components of the δγ_3_δ’ψχ complex, β_2_ and SSB were sequentially assembled on p/t DNA in presence of ADP•AlF_x_ to produce very stable complexes. **h**, Sensorgram showing responses as various components of the δτ_3_δ’ψχ complex, β_2_ and SSB were sequentially assembled on p/t DNA in the presence of ADP•AlF_x_. The biphasic response to δτ_3_δ’ was as expected, since this complex is known also to interact transiently *via* the C-terminal regions of the three τ subunits with ssDNA55, which is present in the 5’ dT_50_ overhang designed for SSB binding. The δτ_3_δ’ bound only *via* this ssDNA quickly dissociated in SPR buffer once protein injection stopped. **i**, Sensorgrams showing δτ_3_δ’ binds more quickly than δγ_3_δ’ to p/t DNA, perhaps due to additional τ−ssDNA contacts that establish promptly55 and pre-concentrate CLC cores near the site of interaction, thereby increasing the efficiency of CLC assembly onto p/t DNA.

**Extended Data Fig. 4.**
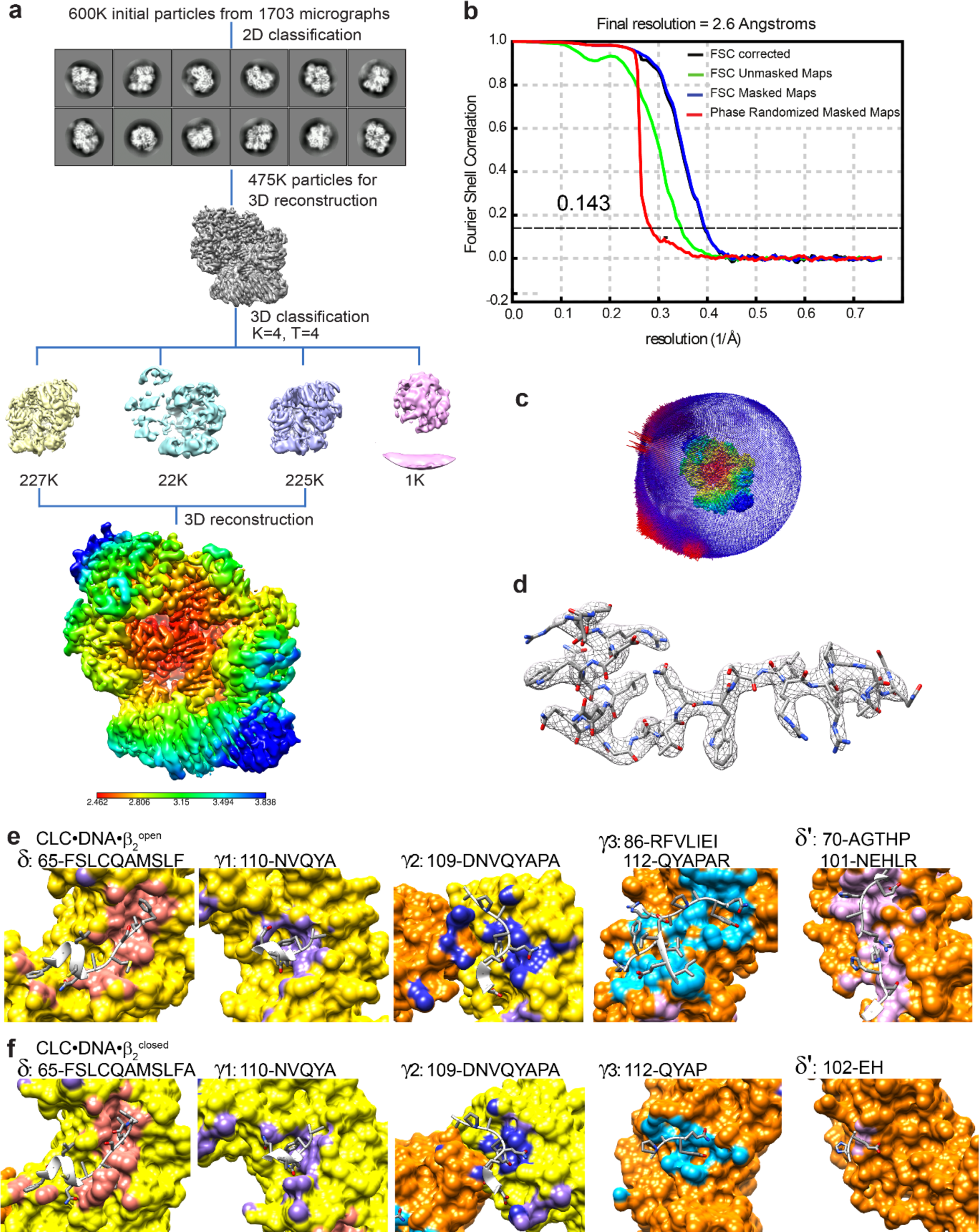
Processing of cryo-EM data of CLC•DNA•β_2_^closed^ complexes containing a p/t DNA with 23-bp dsDNA and a 5’-dT_10_ overhang, related to Fig. 5. **a**, Cryo-EM image processing workflow. **b**, Fourier shell correlation of the final density map. **c**, Angular distribution plot of observed particle orientations. **d**, Unsharpened EM density of the CLC-binding peptide of ψ (Thr2–Glu28), shown as an example of the quality of the map. **e**, **f**, Clamp-binding peptides (ribbons and sticks) in binding pockets of β_2_ of CLC•DNA•β_2_^closed^ and CLC•DNA•β_2_^closed^, respectively. Sequences of peptides involved in binding are shown above the figures.

**Extended Data Fig. 5.**
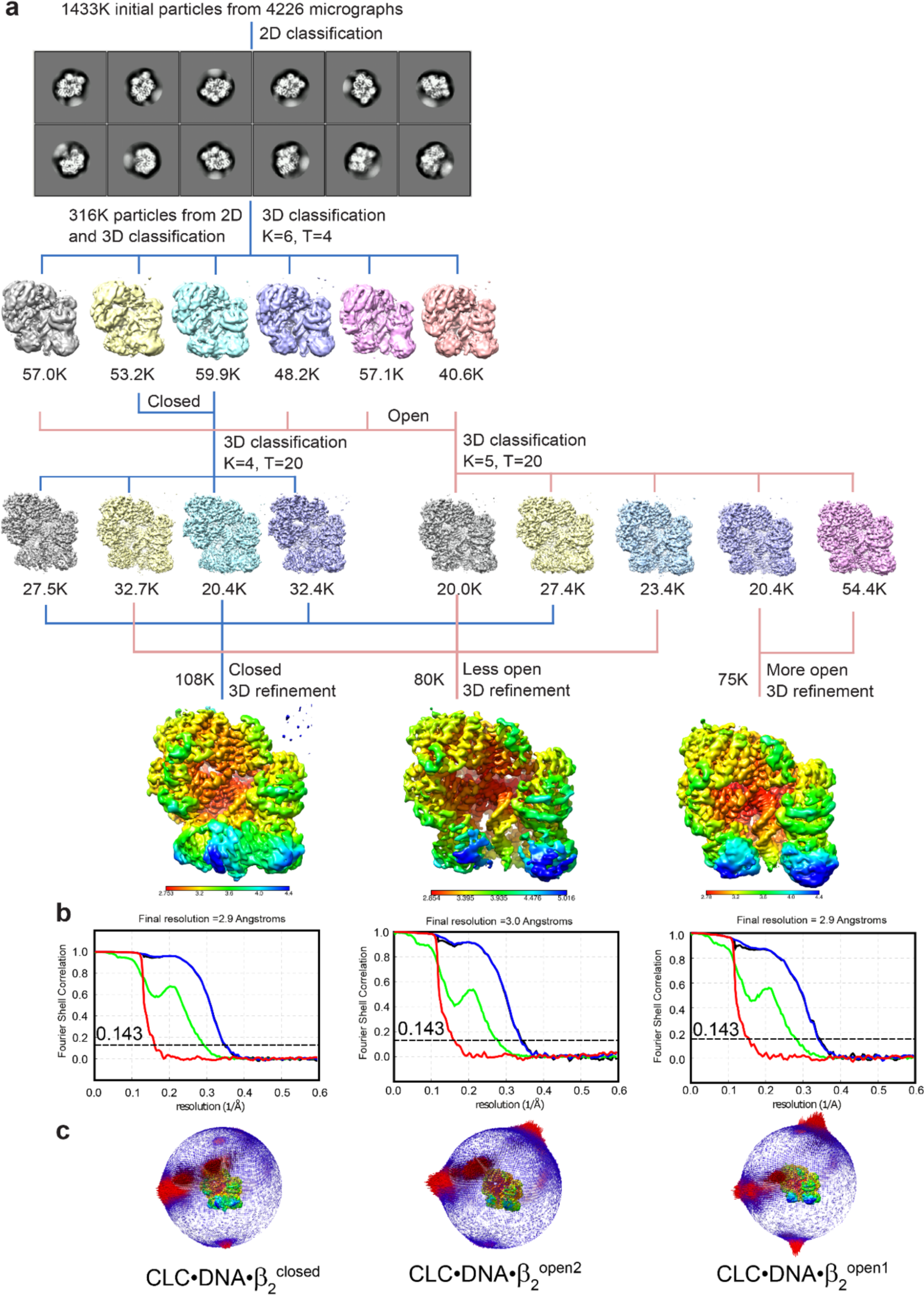
Processing of cryo-EM data of the CLC•DNA•β_2_ complexes containing SSB and a p/t DNA with 23-bp dsDNA and a 5’-dT_45_ overhang, related to Fig. 5. **a**, Cryo-EM image processing workflow. Multiple rounds of 3D classification were performed to separate particles into three distinct classes. Only the last three 3D classification jobs are shown. **b**, Fourier shell correlation of the final density maps. **c**, Angular distribution plot of observed particle orientations.

**Extended Data Fig. 6.**
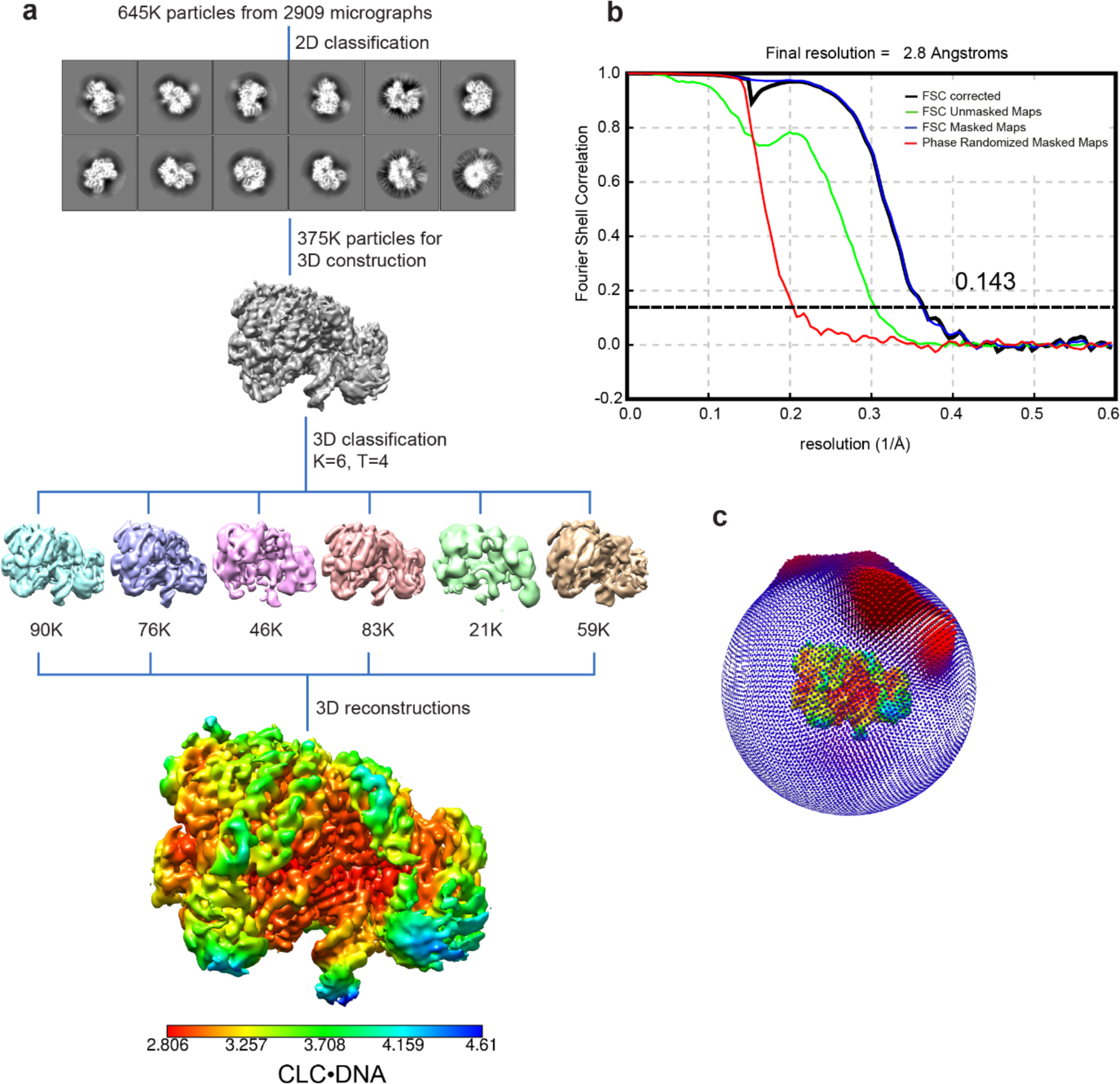
Processing of cryo-EM data of the CLC•DNA complex containing a p/t DNA with 15-bp dsDNA and a 5’-dT_10_ overhang, related to Fig. 5. **a**, Cryo-EM image processing workflow. **b**, Fourier shell correlation of the final density maps. **c**, Angular distribution plot of observed particle orientations.

